# Sensory and Behavioral Components of Neocortical Signal Flow in Discrimination Tasks with Short-term Memory

**DOI:** 10.1101/2020.01.13.904284

**Authors:** Yasir Gallero-Salas, Balazs Laurenczy, Fabian F. Voigt, Ariel Gilad, Fritjof Helmchen

**Affiliations:** Brain Research Institute, University of Zurich, Zurich, Switzerland; Neuroscience Center Zurich, Zurich, Switzerland; Hebrew University Medical School, Department of Medical Neurobiology, Institute of Medical Research Israel-Canada, Jerusalem 9112102, Israel

## Abstract

In neocortex, each sensory modality engages distinct primary and secondary areas that route information further to association areas. Where signal flow may converge for maintaining information in short-term memory and how behavior may influence signal routing remain open questions. Using wide-field calcium imaging, we compared cortex-wide neuronal activity in layer 2/3 for mice trained in auditory and whisker-based tactile discrimination tasks with delayed response. In both tasks, mice were either active or passive during stimulus presentation, engaging in body movements or sitting quietly. Irrespective of behavioral strategy, auditory and tactile stimulation activated spatially segregated subdivisions of posterior parietal cortex (areas A and RL, respectively). In the subsequent delay period, in contrast, behavioral strategy rather than sensory modality determined where short-term memory was located: frontomedially in active trials and posterolaterally in passive trials. Our results suggest behavior-dependent routing of sensory-driven cortical information flow from modality-specific PPC subdivisions to higher association areas.

## INTRODUCTION

Transforming a relevant sensory stimulus into an appropriate action is an operation fundamental to the brain, yet we still understand it poorly. In the neocortex, sensory stimuli of different modalities (e.g., auditory, visual, tactile) are represented in specialized primary and secondary areas. These regions communicate with association areas that in turn route action-instructive signals further towards areas that can hold relevant information in short-term memory and prepare for action (Lyamzin and Benucci, 2019). These transformations require distributed and coordinated activity across many areas. To reveal such large-scale cortical activity patterns, recent advances in wide-field calcium imaging have proven highly beneficial (Allen et al., 2017; Chen et al., 2017; Clancy et al., 2019; Gilad et al., 2018; Makino et al., 2017; Musall et al., 2019; Pinto et al., 2019; Wekselblatt et al., 2016). However, the dependence of neocortical signal flow on specific task requirements (e.g. stimulus modality) and behavioral repertoire (e.g. movement strategy) remains largely unexplored.

A key association area bridging the present (sensory stimulus) to the future (delayed action) is the posterior parietal cortex (PPC), which lies between primary visual and somatosensory areas and projects broadly to areas in frontal and posterior cortex (Harris et al., 2019; Zingg et al., 2014). PPC has been implicated in various functions such as multi-sensory integration (Kuroki et al., 2018; Lippert et al., 2013; Mohan et al., 2018a; Nikbakht et al., 2018; Olcese et al., 2013), decision making (Goard et al., 2016; Pho et al., 2018), and evidence accumulation (Morcos and Harvey, 2016; Odoemene et al., 2018). Given its connectivity and functional role, PPC is a prime candidate to serve as routing area between sensation and short-term memory. The exact anatomical delineation of mouse PPC is, however, still a matter of debate (Glickfeld and Olsen, 2017; Harris et al., 2019; Hovde et al., 2019; Lyamzin and Benucci, 2019; Mohan et al., 2018b; Zingg et al., 2014). A functional delineation of PPC, based on wide-field calcium imaging across distinct behavioral tasks, should provide further insights into its organizational principles.

A particularly intriguing question pertains to short-term memory, the ability of the brain to maintain relevant information in memory over several seconds to guide future actions. Both frontal and posterior cortical areas have been implicated in delay activity related to short-term memory (Goard et al., 2016; Guo et al., 2014; Inagaki et al., 2018; Kamigaki and Dan, 2017; Gilad et al., 2018; Harvey et al., 2012; Morcos and Harvey, 2016; Siegel et al., 2015; see also review by Sreenivasan and D’Esposito, 2019). What determines the routing of cortical signal flow towards these possible locations of short-term memory remains unclear. Several recent studies highlighted the strong influence of behavioral variables, i.e. movement patterns, on cortical dynamics (Clancy et al., 2019; Musall et al., 2019; Salkoff et al., 2019; Stringer et al., 2019). In our own study (Gilad et al., 2018), using a whisker-dependent go/no-go texture discrimination task with delayed response in mice, we found that the location of short-term memory strikingly depends on the movement behavior during sensation. An active strategy, defined as prominent body movements during texture touch, prompted prolonged delay activity in frontomedial secondary motor cortex (M2), likely reflecting a motor plan for licking. In contrast, when mice stayed quiet during the touch, using a passive strategy, delay activity occurred nearly at the opposite cortical pole, posterior and lateral to V1 (which we refer to here as posterolateral association [PLA] areas). These PLA areas presumably held information about a relevant feature of the tactile stimulus. We concluded that behavioral strategy is a key determinant of information flow in this whisker-based tactile task (Gilad et al., 2018; Sreenivasan and D’Esposito, 2019) but it remains unclear whether such behavior-dependent routing generalizes to other tasks based on different sensory modalities.

To address these questions, we here train mice in both auditory and whisker-based tactile discrimination tasks including short-term memory phases. Using wide-field calcium imaging, we uncover a functional subdivision of PPC into sensory modality-specific regions. Furthermore, we find that the location of short-term memory is largely determined by behavioral strategy rather than by the task-relevant sensory modality. Our results emphasize the role of behavior in cortical dynamics and short-term memory, and suggest a critical role of PPC in behavior-dependent routing of neocortical signals to either frontal or posterior high-level cortical areas.

## RESULTS

### Auditory and Tactile Discrimination Tasks with Delayed Response

We trained transgenic mice (expressing GCaMP6f in L2/3 pyramidal neurons) in two go/no-go discrimination tasks with delayed response, using either auditory tones or tactile textures as relevant sensory stimuli (**Figure 1A**). Task design was equivalent except for the sensory modalities of cues and discrimination stimuli. In the auditory task, we trained mice to discriminate between 4-kHz and 8-kHz tones (either serving as go-stimulus). A visual cue signaled the start of each trial, followed by a 2-s long presentation of one of the two tones. In the subsequent delay period mice needed to hold information in short-term memory for several seconds, until a second visual cue signaled that they were allowed to lick for a water droplet as reward in go trials. The tactile task had the same trial structure but instead of tones mice had to discriminate between two textures that were brought in contact with the facial whiskers on the right side of the snout (Chen et al., 2013; Gilad et al., 2018). Either a coarse sandpaper (grit size P100) or a smooth one (P1200) served as go-stimulus. In addition, auditory cues (instead of visual cues as in the auditory task) signaled trial start and the start of the response period. Mice were conditioned to lick for the go-stimulus (‘Hit’ trial; ‘Miss’ if they failed to lick) and to withhold licking for the no-go-stimulus (‘Correct Rejection’, CR; ‘False alarm’ if they erroneously licked). The lick detector was reachable at all times in both tasks. Licks before the response cue (‘Early licks’) were mildly punished with a white-noise sound and a time-out period, as we did for false alarms.

**Figure 1.**
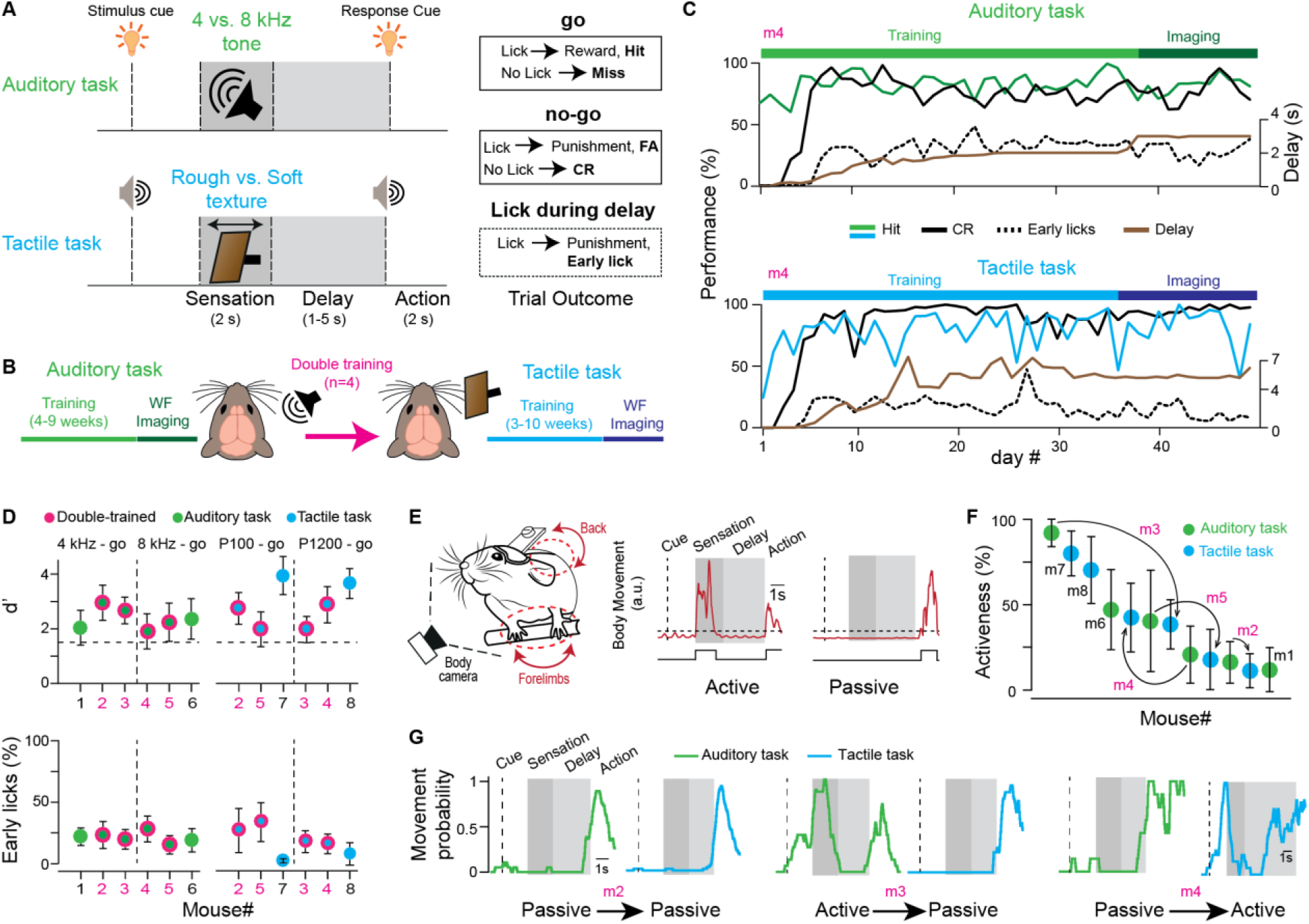
Training, performance, and behavioral strategy of mice in auditory and tactile discrimination tasks. (**A**) Trial structure and possible trial outcomes in the auditory (top) and tactile (bottom) discrimination task with delayed response. (**B**) Timeline of training and wide-field (WF) calcium imaging for the auditory and tactile task. Four mice were sequentially trained in both tasks. (**C**) Performance (Hit and CR rate in percent) of an example mouse (m4) throughout training with increasing delay duration for both tasks. Imaging was performed when the mouse stably performed at expert level with sufficiently long delay. The percentage of Early lick trials is also plotted. (**D**) Performance (d’, top) and fraction of early lick trials (bottom) for each mouse. The respective go-stimuli are indicated on top. Double-trained mice appear twice (pink dots). Dashed line indicates expert threshold at d’ = 1.5. Error bars are SD over expert sessions. (**E**) Left: Video monitoring of body movements during head-fixed behavior. Right: Movements during example active and passive trials (extracted by video analysis). Binary movement vectors (lower traces) were obtained by thresholding (dashed line). Trials were classified as active (left) or passive (right) based on the presence or absence of movements during the sensation period. (**F**) Average movement across imaging sessions for all mice and tasks arranged in descending order. Note how activeness may vary between auditory and tactile task for the double-trained mice (arrows). Error bars are SD over sessions. (**G**) Movement probability calculated from the binary movement vectors for individual example sessions of three double-trained mice. Note the variability of how movement probability may change between tasks.

Mice can learn such discrimination tasks with delayed response over the course of several weeks (Gilad et al., 2018). To compare task-related cortical dynamics directly in the same brain, we trained four mice in both tasks, first the auditory then the tactile task (**Figure 1B**). An additional set of mice was trained in only one task (n = 2 for each task type). For each task, we thus used 6 mice. After initial discrimination learning, we introduced the delay period between sensation and response, which we gradually prolonged during training (**Figure 1C**; total training time 3-10 weeks). Mice learned to withhold licking for several seconds (range 1-4 s for auditory, 1.5-7 s for tactile task), achieving expert-level discrimination performance (d-prime value, d’, above 1.5) while maintaining a relatively low percentage of early licks (23 ± 10% and 20 ± 15% across mice for auditory and tactile task; mean ± s.d.; **Figure 1D**). Once mice had become expert in a task with a sufficiently long delay period, we performed wide-field imaging while animals performed the task.

### Variable Use of Active or Passive Behavioral Strategy across Mice and Tasks

We have previously demonstrated that cortical activity is influenced by the movement behavior of mice during the task trials (Gilad et al., 2018). Specifically, cortical activity is more widespread, especially involving frontal areas, in trials, in which mice actively engage their body during sensory stimulation (e.g., by moving their forelimbs), compared to trials, during which they sit quietly and passively while receiving the stimulus. Hence, it is essential to distinguish between trials representing such ‘active’ and ‘passive’ behavioral strategies, not least because cortical activity at later times, i.e., during short-term memory, turned out to depend on movement behavior as well (Gilad et al., 2018). To discern active and passive trials, we therefore video-recorded body movements while mice solved the two different tasks and extracted trial-related movement vectors (**Figure 1E**). We found that mice adopted variable behaviors, using the active and passive strategy to different degrees on individual trials. We defined ‘activeness’ as the percentage of trials in which an animal used the active strategy. Across all mice, activeness varied widely in both tasks, ranging from 11-92% (**Figure 1F**). Notably, among the double-trained mice some displayed similar activeness across tasks whereas others changed their preferential use of either the active or the passive strategy. For example, mouse 3 and 5 substantially reduced their overall activeness in the tactile compared to the auditory task while mouse 4 increased its activeness. On the contrary, mouse 2 maintained its preferred use of the passive strategy throughout both tasks (**Figure 1G**). We conclude that individual mice adopt a particular behavioral repertoire to solve each task, characterized by preferential use of either the active or the passive strategy, but that this repertoire may flexibly change from task to task.

### Activation of Distinct PPC Subdivisions in the Auditory and Tactile Task

How does trial-related cortical activity differ between the two tasks? To simultaneously monitor all areas across dorsal cortex, we used wide-field calcium imaging through the intact skull above the left hemisphere (Gilad et al., 2018; Vanni and Murphy, 2014) as our mice were triple transgenic mice expressing GCaMP6f in L2/3 pyramidal neurons (Madisen et al., 2015) (**Figure 2A**; Methods). To localize primary sensory areas and register brains to a reference atlas, each mouse underwent a sensory mapping session under anesthesia (**Figure 2A**; Methods). By presenting different stimuli, we localized barrel cortex (BC), forelimb and hindlimb cortices (FL and HL), visual cortex (V1) and primary auditory cortex (A1). Taken these locations as anchors, together with anatomical landmarks (i.e. bregma and lambda), we further aligned each brain to the Allen Mouse Common Coordinate Framework (**Figure S1**; Methods; Oh et al., 2014). This registration allowed us to identify corresponding areas across mice and pool the respective calcium signals.

**Figure 2.**
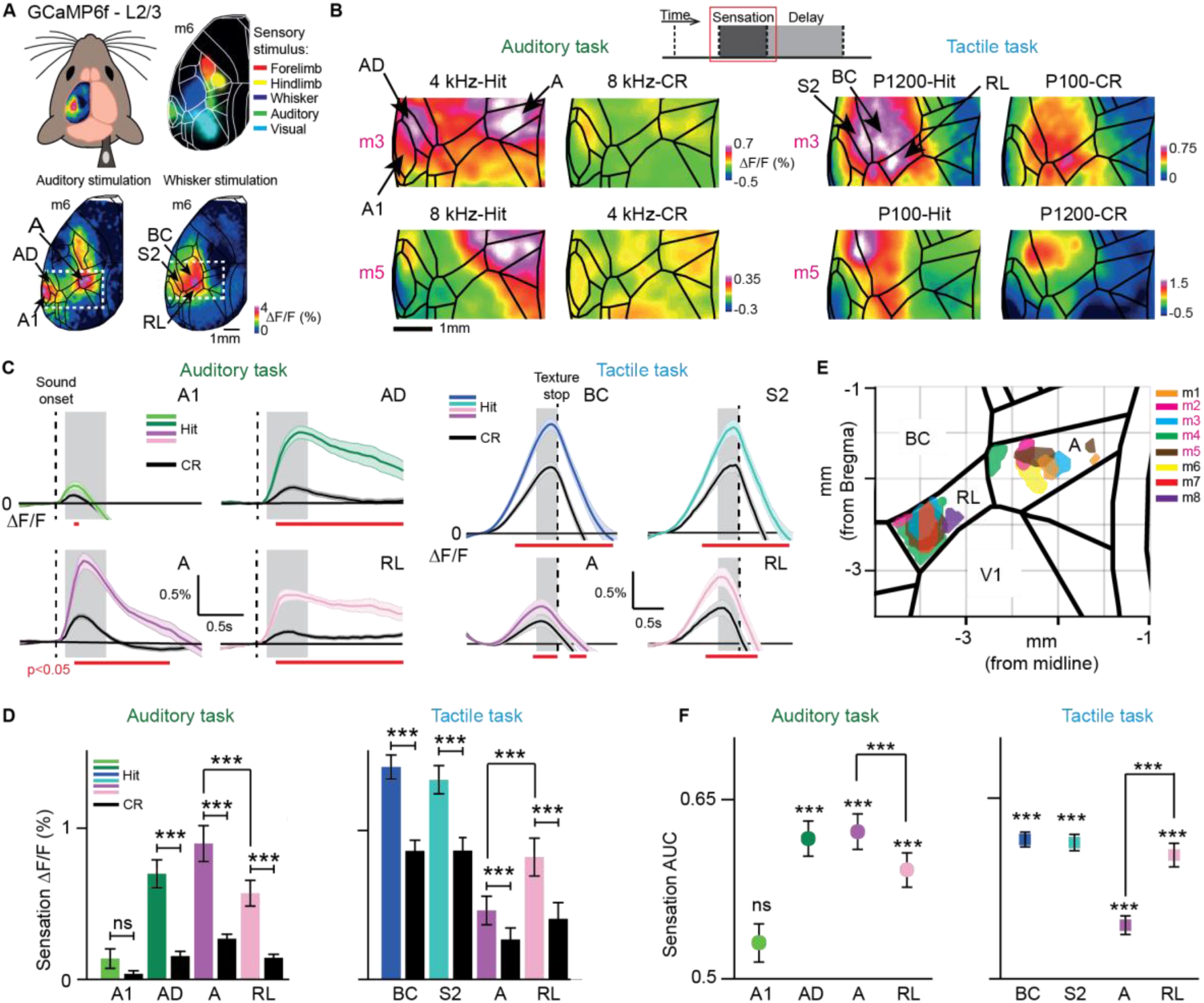
Modality-specific activation of cortical areas during sensation, including distinct subdivisions of PPC. (**A**) Top-left: Wide-field calcium imaging across the left hemisphere. Top-right: Merged sensory-evoked activity maps for registration to the Allen atlas. Bottom: Example single-trial activity maps in response to auditory (left) and whisker (right) stimulation during anesthesia.(**B**) Example sensation maps of two double-trained mice for Hit and CR trials in both tasks (session-averages including active and passive trials). Zoom-in corresponds to dashed white boxes in (A). Maps were calculated from early time periods during sensation (gray boxes in c). Color scale bars indicate minimum and maximum ΔF/F.(**C**) Average ΔF/F time course for Hit versus CR in A1, AD, A and RL in the auditory task (left) and in BC, S2, A and RL in the tactile task (right). Error bars are SEM across sessions. Dashed line indicates sound onset (left) and texture stop (right; first touches typically occurred 0.5-1 s before texture stop). Gray boxes indicate time windows for calculation of sensation maps. Red horizontal lines indicate time periods of significant Hit vs. CR difference (p < 0.05; One-way ANOVA, Least Significant Difference corrected).(**D**) Mean sensory-evoked ΔF/F changes in early time windows (gray boxes in c) for Hit versus CR trials in A1, AD, A and RL in the auditory task (left, n = 84 sessions from 6 mice) and in BC, S2, A and RL in the tactile task (right, n = 78 sessions from 6 mice). Error bars are SEM across sessions.(**E**) Location of the 10% most active pixels within RL and A for each animal in both tasks.(**F**) Hit versus CR discrimination power, calculate as area under the ROC curve (AUC) for A1, AD, A and RL in the auditory task (left, n = 84 sessions) and for BC, S2, A and RL in the tactile task (right, n = 78 sessions from 6 mice). We calculated significance of discrimination power for each area by comparing with shuffled trials. Error bars are SEM across sessions.*p < 0.05, **p < 0.01, ***p < 0.001; n.s., not significant; Wilcoxon signed-rank test. See also Figures S2-3.

We first analyzed spatiotemporal cortical activity upon sensory stimulation by creating spatial activity maps (**Figure 2B**, grouping together active and passive trials) and by extracting ΔF/F time courses for sensory-related cortical areas (**Figure 2C**; see **Figure S2** for all areas). In the auditory task, we observed tone-related activity changes in primary auditory (A1), auditory dorsal (AD) and auditory posterior (AP) cortices, with AD showing the highest activation level for the go-tone and significant discrimination between Hit and CR trials (irrespective of go-tone type, **Figure 2B-D**; p < 0.001; Wilcoxon signed-rank test). Interestingly, tone-evoked activity in A1 was highly variable between mice, with some animals displaying decreases rather than increases, and averaged across animals did not significantly discriminate Hit and CRs (**Figures S2 and S3**; see also Discussion). In the tactile task, touch-evoked activity was strong in BC and secondary somatosensory cortex (S2), being significantly higher in Hit versus CR trials, irrespective of go-texture type (**Figure 2B-D; Figure S2;** p < 0.001; Wilcoxon signed-rank test).

In addition to the relevant primary and secondary cortices, areas representing PPC showed strong stimulus-evoked activity. In the auditory task, tone stimulation most strongly activated area A in the medial part of PPC. In contrast, the rostrolateral area RL, as lateral part of PPC, was significantly more engaged in the tactile task (**Figure 2B-D**; p < 0.001; Wilcoxon signed-rank test). The spatial separation and differential engagement of these two areas is clearly visible when plotting the location of the 10% most active pixels during sensation in A and RL for each animal in the auditory and tactile tasks, respectively (**Figure 2E**). For the four double-trained mice, the distance between activation peaks in A and RL was 1.7 ± 0.25 mm. We conclude that PPC does not function as a single integrative hub but that different sensory modalities engage spatially segregated subdivisions of PPC (as unambiguously demonstrated in the double-trained mice).

To investigate how well each cortical area could discriminate between Hit and CR trials, we performed receiver operating characteristics (ROC) of sensory-evoked ΔF/F amplitudes across trials and calculated the area under the curve (AUC) (**Figure 2F**; AUC = 0.5 indicates chance-level discrimination and AUC values closer to one indicate high discrimination power). Pooled across mice, AD but not A1 could discriminate significantly above chance level in the auditory task. Within PPC, area A had significantly higher discrimination power than RL (p < 0.001; Wilcoxon signed-rank test). In the tactile task, whisker-related areas (BC, S2) and RL showed high Hit/CR discrimination, with RL discriminating significantly better than area A (p < 0.001; Wilcoxon signed-rank test).

We also compared sensory-evoked responses for the different behavioral strategies. In general, cortical activity in the sensation period was more widespread in active compared to passive trials, engaging further somatosensory areas as well as motor-related frontal areas (**Figure S4**). In active trials, sensory-evoked responses were enhanced in the relevant primary, secondary, and PPC areas for each task type and showed Hit/CR discrimination power comparable to passive trials (**Figure S4**). Notably, the preferential activation of A and RL in the auditory and tactile task, respectively, was present in both active and passive trials. The behavior-dependence of cortical activity during sensory integration, with stronger and more widespread activity in active trials, also raises the question in how far behavioral strategy may influence signal flow in the subsequent delay period.

### Location of Short-Term Memory Depends on Behavioral Strategy but not Sensory Modality

We therefore next analyzed the delay period, during which mice had to maintain information in short-term memory. We treated active and passive trials separately and analyzed only trials, in which at least the first second of the delay period was free of movement (Gilad et al., 2018). In addition, we truncated the delay period for each trial when the first movement occurred (anticipatory movements preparing for licking). The delay activity maps reported here thus include only periods, in which mice were sitting quietly. As an example, we plot in **Figure 3A** the activeness of all recorded sessions for the double-trained mouse #3 and exemplify single-trial delay maps for active and passive Hit trials from example sessions of each task (**Figure 3B**). Delay maps were highly distinct for active compared to passive trials, both for the tactile task, consistent with our previous study (Gilad et al., 2018), and for the auditory task. Across the two tasks, delay activity maps were similar for trials of the same strategy, showing highest activity in M2 near the midline for active trials and in PLA areas for passive trials (**Figure 3B and 3C**; see **Figure S5** for a detailed analysis). PLA areas mainly comprised areas LM, LI, PL and POR.

**Figure 3.**
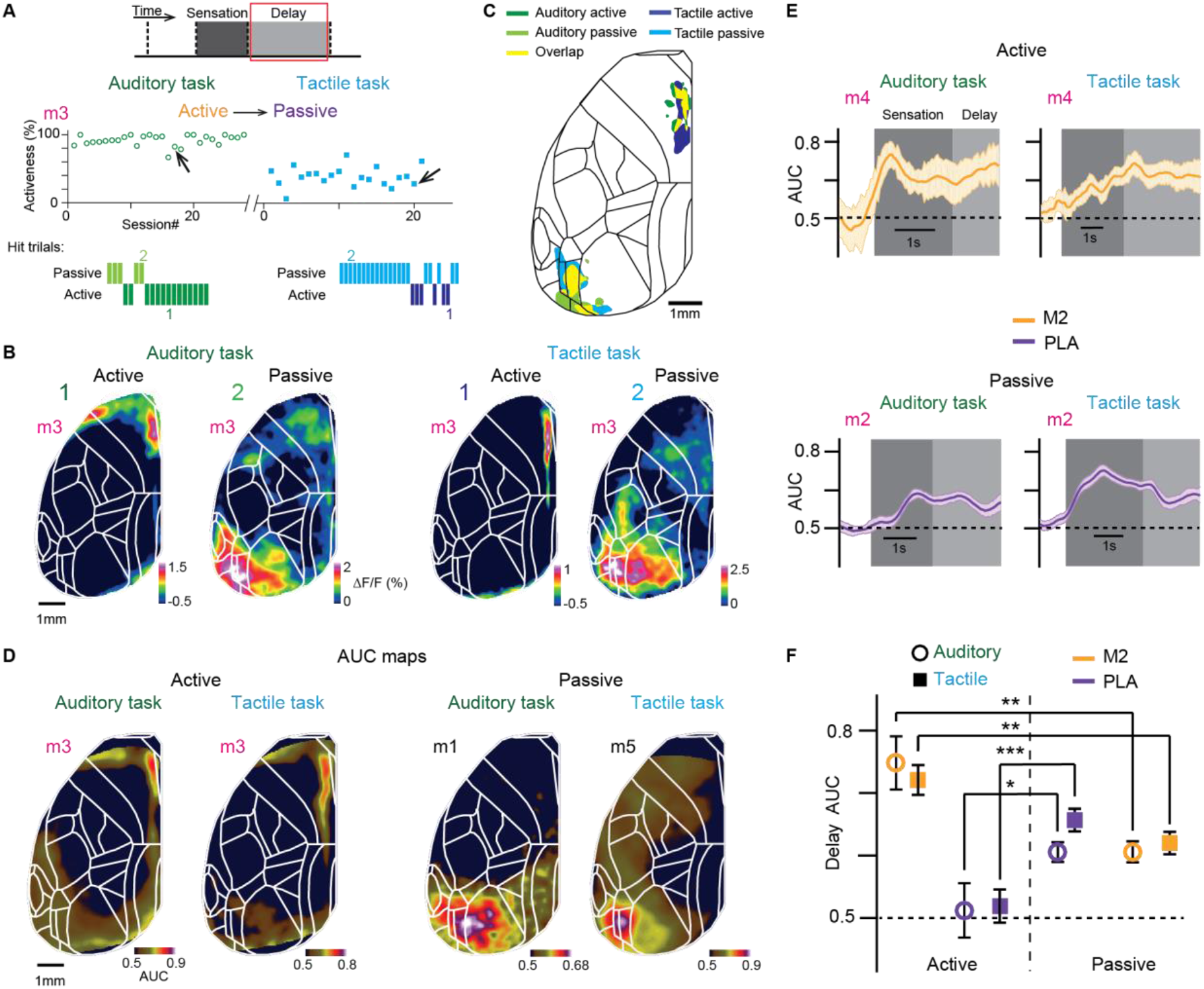
Location of delay activity with high Hit/CR discrimination power differs for active and passive behavior. (**A**) Top: Activeness across imaged session in a mouse (mouse 3) trained in the auditory (left) and tactile (right) tasks. Low: Hit and passive trials within an example session indicated by an arrow. (**B**) Example single-trial active (1) and passive (2) delay maps in the auditory (left) and tactile (right) tasks. Color scale bar indicates ΔF/F percentage. (**C**) Location of the 10% most Hit/CR discriminative pixels in frontomedial M2 (active trials) and in PLA areas (passive trials) in both tasks (obtained from example sessions from each mice). Yellow areas indicate overlap across tasks. (**D**) Example session delay AUC maps (for Hit/CR discrimination) for both auditory and tactile task for active (left) and passive (right) strategy. Color scale bar indicates AUC. (**E**) Average AUC time course for Hit/CR discrimination in M2 for active trials and in PLA areas for passive trials in both tasks. Error bars are SEM across sessions. (**F**) Average delay AUC for Hit/CR discrimination in M2 and PLA areas for the active and passive strategy in both tasks. Error bars are SEM across sessions (auditory task: n = 70 passive sessions from 6 mice, n = 28 active sessions from 4 mice; tactile task: n = 64 passive sessions from 6 mice, n = 30 active sessions from 6 mice). *p < 0.05, **p < 0.01, ***p < 0.001; n.s., not significant; Wilcoxon signed-rank test. See also Figures S5 and S6.

To analyze Hit/CR discrimination power during the delay period, we calculated the AUC of ROC for each pixel and plotted delay AUC maps (**Figure 3D**). We also extracted AUC time courses throughout the trial (**Figure 3E**). We found that M2 displayed significantly higher discrimination power during the delay period in active compared to passive trials (**Figure 3F**; p<0.01; Wilcoxon signed-rank test). Conversely, PLA areas showed significantly better discrimination for passive compared to active trials in both tasks (p<0.05; Wilcoxon signed-rank test). In passive trials, M2 also exhibited above-chance discrimination power although ΔF/F activity for this frontomedial area on average was much smaller than in active trials (**Figure 3F**). These results suggest that the location of persistent activity during short-term memory is determined predominantly by behavioral strategy, irrespective of task modality.

### First Sensory Modality then Behavioral Strategy Governs Cortical Dynamics within Trials

To further quantify the impacts of sensory modality and behavioral strategy on cortical activity, we defined two indices based on how well cortical activity discriminated between either auditory and whisker modalities (‘task index’) or between active and passive strategies (‘strategy index’) (**Figure 4A**; Methods). We calculated these indices for each imaging frame during the trial period separately, focusing on the Hit trials of the four double-trained mice so that we could directly compare the same cortical areas across all conditions. To create index maps, we averaged the index values for each pixel over either the early sensation period or the delay period. The resulting maps for task index and strategy index for the sensation period confirmed high discrimination power of sensory-related areas upon sensory stimulation but low discrimination of behavioral strategies across all cortical areas (**Figure 4B**). During the delay period, on the contrary, M2 and PLA areas showed high discrimination between active and passive strategy whereas most of cortex showed low discrimination between tasks (**Figure 4C**).

**Figure 4.**
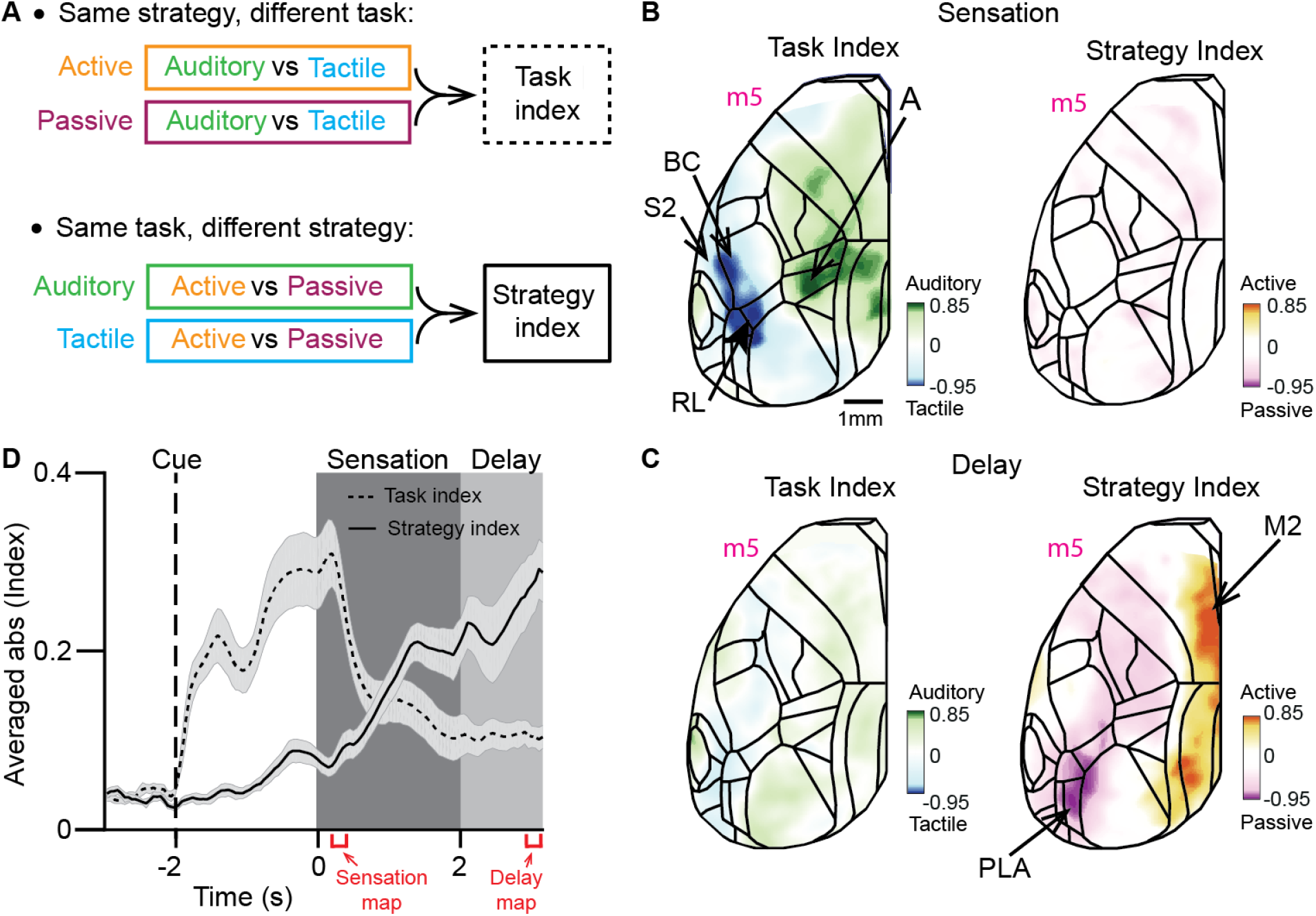
Task modality and behavioral strategy dominate cortical dynamics during sensation and delay, respectively. (**A**) Schematic illustrating the calculation of task index (top, merging the result for the two strategies) and strategy index (bottom, merging the results for the two tasks). (**B**) Task index map (left) and strategy index map (right) during early sensation (left red bracket in D, 0.1 to 0.2 s) for one example mouse. Color scale bars indicate range of index values (−1, 1). Green scale indicates pixels with high discrimination power for auditory task; blue scale indicates pixels with high discrimination power for tactile task. (**C**) Task index map (left) and strategy index map (right) during the delay period (right red bracket in D, 2.7 to 3 s) for the same example mouse as in B. Color scale bars indicate strategy index (−1 to 1). Orange scale indicates pixels with high discrimination power for active strategy; purple scale indicates pixels with high discrimination power for passive strategy. For both indices, zero indicates absence of discrimination power. (**D**) Average time course of the absolute value of task and strategy indices (0-1). Error bars are SEM across brain regions (n=26 brain regions, Figure S1).

Finally, to evaluate the temporal progression of cortical dynamics we averaged the absolute value of task and strategy indexes across areas for each imaging frame within the trial period (**Figure 4D**). The average task index increased after the initial trial start cue, peaked during early sensation, and then decreased towards the delay period. Conversely, the strategy index remained low during early sensation but increased towards the end of sensation and reached the highest level during the delay period. This analysis confirms and directly illustrates that large-scale cortical dynamics are dominated by the modality of the relevant external sensory stimulus early during the task trials but then – after the stimulus has been received and information has to be maintained in short-term memory – is predominately governed by internally produced behavior. Apparently, neocortical signals are differentially routed to either frontomedial or posterolateral areas in a behavior-dependent manner to hold decisive information.

### Anatomical Connectivity Supports the Activity Maps Observed for Sensation and Short-Term Memory

Given the sensory modality-dependent activation during sensation and the differential signal flow towards the delay period, we asked whether the anatomical connectivity between task-specific sensory areas supports the observed signal flow patterns. To this end, we downloaded projection data from the Allen Mouse Brain Connectivity Atlas (Harris et al., 2019) for all sensory-related areas. Then, we created a connectivity matrix for the sensory-related areas relevant in our tasks, averaging across all the available experiments in Cre lines for each area. In agreement with our functional activity maps, projections among modality-specific areas (A1, AD, A versus BC, S2, RL) are generally stronger than across modalities (**Figure 5A**). The strongest connections occur between primary and secondary sensory cortices (reciprocal, A1↔AD and BC↔S2) and PPC subdivisions (reciprocal, A↔RL). Additionally, area A projects more strongly to AD than to A1, suggesting that communication between A1 and A occurs principally through AD (Zhong et al., 2019) (**Figure 5B**). On the other hand, RL strongly projects to BC and S2 with weaker reciprocal connections (**Figure 5B**). In summary, we find a modality-specific anatomical substrate that may underlie the observed activation patterns during sensory integration.

**Figure 5.**
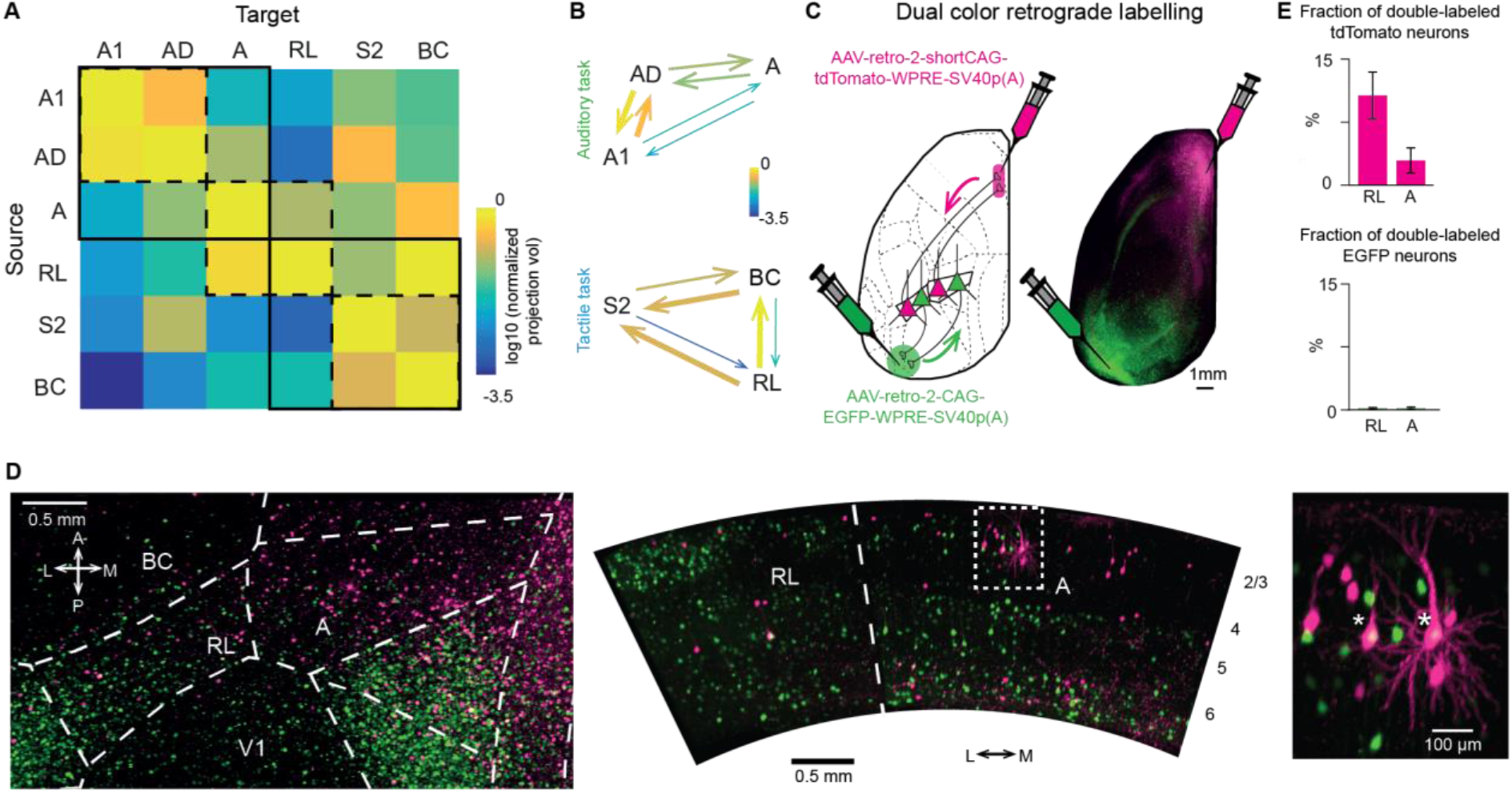
Anatomical connections support functional segregation. (**A**) Average normalized projection volume from sensory related areas in both tasks downloaded from the Allen Mouse Brain Connectivity Atlas. Solid and dash squares suggest groups of auditory, somatosensory and parietal areas. (**B**) Sensory related areas proposed dynamics during sensation for auditory (top) and tactile (bottom) tasks. Arrow thickness reflects the connection strength. (**C**) Retrograde labelling of neurons projecting to M2 and PLA areas by inducing tdTomato and EGFP expression, respectively. Arrows indicate retrograde labeling. (**D**) Light-sheet microscopy images of areas RL and A in a cleared mouse brain in dorsal (left) and coronal (middle) view. A higher magnification view of the box indicated in the middle is shown on the right. Asterisk denotes double-labelled cells. Image histogram was independently adjusted in RL and A in the coronal view. The tissue clearing process induced an expansion of the tissue of approximately 1.5x. (**E**) Quantification of the fraction of M2-projecting neurons in RL and A that also project to PLA areas (top) and of PLA-projecting neurons in RL and A that also project to M2 (bottom). Error bars are SEM across mice (n=3).

Regarding the distinct delay activity patterns, previous studies have already established that both RL and A project to frontal and posterolateral areas (Harris et al., 2019; Oh et al., 2014; Wang et al., 2012). However, it remains unknown whether these pathways originate from segregated neuronal subpopulations in the PPC areas or from the same pool of projection neurons. To investigate whether individual neurons in RL and A project to both frontal and posterior areas or only to one of these, we injected 3 mice with retrograde viruses coding for two differently colored fluorescent proteins in M2 and PLA areas (AAV-retro-2-shortCAG-tdTomato-WPRE-SV40p(A) and AAV-retro-2-CAG-EGFP-WPRE-SV40p(A), respectively) (**Figure 5C**). After 4 weeks, we cleared the brains using a CLARITY protocol (Chung et al., 2013) and performed whole-brain imaging with our mesoSPIM light-sheet microscope (Voigt *et al*., 2019). We found both PLA-projecting (EGFP-expressing) as well as M2-projecting (tdTomato-expressing) neurons in RL and A (**Figure 5D**). Only few of the identified projection neurons were double-labelled. Quantification revealed that 10% or less of M2-projecting neurons in RL and A also project to PLA areas and negligible co-expression was found in PLA-projecting neurons (**Figure 5E**).

These findings demonstrate that the output projections from PPC (both in A and RL) originate from largely segregated pools of projection neurons, which could be an anatomical substrate of the observed differential routing of information to either M2 or PLA areas for short-term memory.

## DISCUSSION

We have shown that sensory modality and behavioral strategy are determining factors of signal flow through neocortical association areas during sensation and short-term memory. Below we discuss the distinct activation patterns that we observed during the sensation period, especially for PPC, and the behavior-dependent location of short-term memory, which we found to generalize across sensory modalities. We propose a working model for cortical signal routing for the go/no-go type of sensory discrimination tasks investigated here, which is consistent with anatomical connectivity. We conclude that considering trial-by-trial variations in behavior is essential when analyzing cortical signal flow, especially for conditions engaging short-term memory circuits for maintenance of relevant information.

### Particularities of Auditory Evoked Cortical Signal Flow

Compared to passive listening, sound-evoked activity in A1 may be enhanced or reduced in different auditory tasks (Francis et al., 2018; Jaramillo and Zador, 2011; Kato et al., 2015; Kuchibhotla et al., 2017; Otazu et al., 2009; Xin et al., 2019). Moreover, the causal involvement of A1 in auditory tasks remains controversial in view of conflicting results depending on task and inactivation method (Ohl *et al*., 1999; Jaramillo and Zador, 2011; Gimenez, Lorenc and Jaramillo, 2015; Kato, Gillet and Isaacson, 2015; Kuchibhotla *et al*., 2017; Talwar, Musial and Gerstein, 2017; Xin *et al*., 2019). For example, ablations studies in rodents suggest that A1 is not required for pure tone discriminations (Gimenez et al., 2015; Ohl et al., 1999), whereas recordings and lesions studies in monkeys indicate A1 involvement (Colombo et al., 1990; Sakurai, 1994). Here, we found small and variable tone-evoked responses in A1 of mice during task performance. Instead, AD showed the highest activity and Hit/CR discrimination power during sensation (**Figure S3**; see Methods for delineation of A1-AD-S2 boundaries). These findings cannot be explained by damage to A1 or a lack of GCaMP6f expression since we clearly observed A1 activation in all mice during sensory mapping (**Figure S1**) as well as upon task replay during anesthesia and in response to auditory cues in the tactile task (**Figure S3**). Rather, they indicate a particularly strong role of higher auditory areas during task engagement, in line with recent reports from various mammalian species (Atiani et al., 2014; Dong et al., 2013; Elgueda et al., 2019; Niwa et al., 2013; Tsunada et al., 2015). The strong connections of AD with PPC and secondary motor cortex (Harris et al., 2019; Zhong et al., 2019) might also indicate a more prominent role compared to A1 in auditory decision-making tasks, a notion that remains to be further investigated and causally tested in the future.

### Functional Organization of PPC with Respect to Sensory Modalities

Pioneering work on mouse PPC activity during visually-guided navigation tasks focused on the area at the border between A and AM (Harvey et al., 2012). Presumably influenced by this work, subsequent studies targeted this medial part of PPC, too, regardless of which sensory modality was used (Goard et al., 2016; Guo et al., 2014; Le Merre et al., 2018; Zhong et al., 2019). Connectivity studies suggest, however, a mediolateral organization of connections between PPC and distinct sensory areas (Harris et al., 2019; Lee et al., 2011; Wang et al., 2012; Wilber et al., 2015; Zingg et al., 2014). In fact, somatosensory responses have been previously reported in area RL (Gilad et al., 2018; Mohajerani et al., 2013; Mohan et al., 2018a; Olcese et al., 2013). By training the same mouse in auditory and whisker-based discrimination tasks, we here found a clear sensory modality-dependent recruitment of PPC subdivisions along the mediolateral axis. Anatomy supports this finding with A1-AD-A and BC-S2-RL forming tightly connected triangles. Recent electrophysiological recordings in the rat point to an even finer graded somatotopy representing the whisker rows in RL (Mohan et al., 2019). Interestingly, another recent study revealed that RL, which is typically considered a higher order visual area, is specialized for encoding visual stimuli very close to the mouse, within reach of the whiskers, suggesting the existence of a visuo-tactile map of near space in RL (La Chioma et al., 2019). While these insights provide some clarification of the gross functional organization of rodent PPC, further work is needed to disentangle the partially overlapping and intermingled connections with visual, auditory, and somatosensory areas that likely form the basis of the multi-sensory integrative power of PPC.

### Variable Behavioral Strategies for Solving Sensory Discrimination Tasks

Our study confirms and extends our previous finding that mice can use either an active or a passive approach during stimulus presentation to solve a sensory discrimination task (Gilad et al., 2018). In active trials, defined by clear body movements such as limb movements, body stretching and vigorous whisking, we observed widespread activity across cortex in addition to the specific stimulus-evoked responses. This behavior-related activity likely reflects activity involved in movement execution and control, proprioceptive signals, and activity evoked by body parts touching external objects. This behavior-related component represents a substantial part of cortex-wide activity, in line with recent studies highlighting the strong influence of behavioral variables on cortical activity (Clancy et al., 2019; Musall et al., 2019; Salkoff et al., 2019; Stringer et al., 2019). Does higher activeness lead to better performance? Generally, it is important to emphasize that mice can reach high performance levels with either strategy. Nonetheless, for the tactile task, in which mice can engage their body and actively whisk in anticipation of the texture arrival to enforce the whisker-texture touch, activeness indeed positively correlates with performance (Gilad et al., 2018) (**Figure S7**). In this regard, it was surprising for us to find a similar range of activeness in mice for the auditory task, given that no active process contributing to auditory sensing is obvious (Schroeder et al., 2010; not considering echolocation and head and pinnae movements for sound localization). Different from the tactile task, however, activeness did not correlate with performance in the auditory task (**Figure S7**). So why would a mouse use the active strategy to solve the auditory task? One explanation could be that the active strategy helps to prevent uninstructed licks during the delay period, consistent with the negative correlation between activeness and percentage of early licks (**Figure S7**). An alternative explanation could be that mice regulate their motor variability during reinforcement learning and settle on a movement pattern that appears to them to lead to, and thus may be necessary for, successful outcome (Dhawale et al., 2019). This explanation is less compelling, though, for the mice that flexibly use both active and passive strategies. This interpretation would explain, however, our observation that individual mice do not necessarily show the same activeness in the two consecutively trained tasks. In any case, our data highlight the broader relevance of distinguishing active and passive behavioral trials as they may support similar task performance while significantly modulating cortical activity.

### Behavioral Strategy Guides Signal Flow to Distinct Short-term Memory Locations

The recruitment of cortical areas for short-term memory depends on information extracted during sensation (Gilad et al., 2018; Lee et al., 2013; Sreenivasan and D’Esposito, 2019). Differential recruitment may be implicitly instructed by task demands (Lee et al., 2013) or may be influenced by individual preferences such as behavioral strategy (Gilad *et al*., 2018). Here, by training the same mouse in two tasks (auditory and tactile), we showed that the location of persistent delay activity during a short-term memory phase is determined by strategy rather than sensory modality (**Figure 4**). This suggests that information extracted during sensation depends on internal goals (what information the animal choses to remember) irrespective of task sensory modality. In fact, M2 is known to transform multisensory information into an adequate motor plan (reviewed by Barthas and Kwan, 2017) as well as to encode choice (Sul et al., 2011) and display high activity during a delay period (Gilad et al., 2018; Murakami et al., 2014). Therefore, we speculate that, in active trials, both auditory tones and textures are transformed into a motor plan (go or no-go), which is maintained in M2 during the delay period. This immediate shift to frontal cortex after sensation is possible for the go/no-go paradigm. In passive trials, however, we propose that higher-level (abstract) features of texture and sound, for example roughness and pitch, stimulus identity (Gilad et al., 2018; Lee et al., 2013) or value (Ramesh et al., 2018; Shuler and Bear, 2006) are extracted during sensation and that this information is maintained in PLA areas. In humans, it is well known that multisensory information reaches temporal areas (Beauchamp, 2005)—a potential homologue of the mouse PLA areas (Wang et al., 2012)—during sensation of objects (Amedi et al., 2005; Lucan et al., 2010) and during sensory working memory (Quak et al., 2015). Since PLA areas project to retrohippocampal areas, as well as parietal and temporal cortices (Harris et al., 2019; Wang et al., 2012), they could be involved in retrieving long-term memory for matching of the stimulus presented with a stored template. These hypothesis remain to be tested using behavioral paradigms that instruct what exact information must be maintained during the delay period (Esmaeili and Diamond, 2019; Lee et al., 2013; Liu et al., 2014).

### PPC as Signal Router in Neocortex

How is information differentially routed to either frontal or posterior cortical regions depending on behavioral strategy? We hypothesize that a candidate area must 1) be activated during sensation and 2) project to both M2 and PLA areas (Gilad et al., 2018). Our own data and previous data indicate that RL and A fit these criteria (Harris et al., 2019; Wang et al., 2012). In particular, we found that the projection neurons in RL and A project almost exclusively to either frontal or posterior areas, suggestion a pathway segregation that could explain the observed dichotomy of cortical activation patterns in the delay period. These areas seem capable of routing information in a strategy-dependent manner. In active trials, we speculate that movements during sensation may facilitate shifts of activity towards frontal areas, for example through feedback from motor areas that might bias (pre-depolarize) anterior-projecting PPC neurons, effectively lowering their threshold for activation (**Figure 6**).

**Figure 6.**
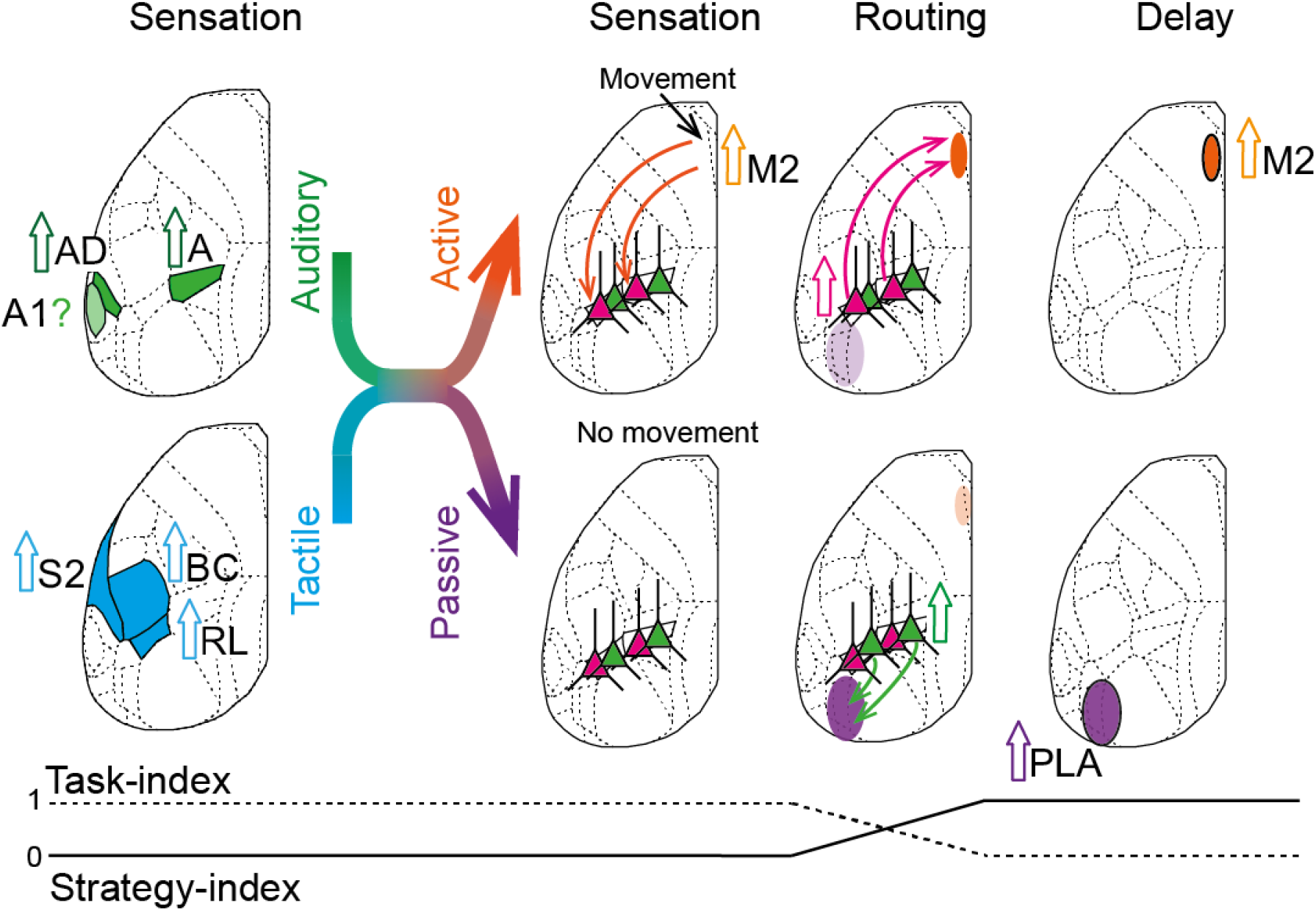
Working model of cortical dynamics during sensation and delay periods. During sensation, taskdependent areas are engaged, regardless of strategy. In the active strategy, motor areas active during sensation (e.g M2) may facilitate the routing of information towards frontal areas in both tasks (e.g. by pre-depolarizing anterior-projecting PPC neurons). In the absence of movement (passive strategy), information flows towards posterior cortices through posterior-projecting PPC neurons. The switch from high task-index to high strategy-index is depicted at the bottom.

In this scenario, movements would promote the transformation of the stimulus received into an adequate motor plan (Barthas and Kwan, 2017). Alternatively, movements could emerge as a side effect of frontal activation during motor planning. In passive trials, the absence of major movements and low activity in frontal areas would make routing of information towards frontal areas less likely. In this case, posterior-projecting PPC neurons might be more easily excited so that information would be routed towards posterolateral association areas. In this way, PPC could act as a major routing area for controlling signal flow across cortex, influenced by both external stimuli as well as behavioral state. Future work employing pathway-specific recordings and manipulations of neuronal subpopulations should help to further substantiate this presumed routing function of PPC.

## AUTHOR CONTRIBUTION

Y.G-S., A.G. and F.H. designed the experiments; Y.G-S., and A.G. conducted the experiments; B.L. designed the auditory behavioral apparatus and wrote analysis code; F.F.V. performed light-sheet imaging on cleared brains; Y.G-S, A.G. and F.H. analyzed the data and wrote the paper.

## ACKNOWLEDGEMENT

We thank Ladan Egolf for managing the transgenic mouse lines, Dubravka Göckeritz-Dujmovic for clearing of brain tissue, Philipp Bethge for help with light-sheet imaging on cleared brains and Erik Fagerholm for help with the task indexes equations. We thank Shankar Sachidhanandam, Benjamin Grewe, Christopher Lewis, and Dayra Lorenzo for comments on the manuscript. This work was supported by grants from the Swiss National Science Foundation (310030B-170269; F.H.; Sinergia project CRSII5_180316; F.H.), and the European Research Council (ERC Advanced Grant BRAINCOMPATH, project 670757; F.H.).

## METHODS

All experimental procedures were carried out according to the guidelines of the Veterinary Office of Switzerland and following approval by the Cantonal Veterinary Office in Zurich (licenses 285/2014, 211/2018).

### Mice and surgical procedures

Eight adult male mice were included in this study, all of which were triple transgenic Rasgrf2-2A-dCre;CamK2a-tTA;TITL-GCaMP6f mice expressing GCaMP6f in layer 2/3 pyramidal neurons of the neocortex. This intersectional genetic strategy allows for specific yet high expression of GCaMP6f (Madisen et al., 2015). Because this line expresses a destabilized Cre (dCre), it requires stabilization by trimethoprim (TMP) in order to express the indicator. We induced GCaMP6f expression in each mouse as follows: TMP (Sigma T7883) was reconstituted in Dimethyl sulfoxide (DMSO, Sigma 34869) at a saturation level of 100 mg/ml and intraperitoneally injected (150 μg TMP/g body weight; 29g needle) at least one week before imaging commenced (typically before training onset).

To perform wide-field calcium imaging chronically (over several months) we used the minimally invasive intact skull preparation originally described by Silasi and colleagues (Silasi et al., 2016). We followed the procedures as described previously (Gilad et al., 2018). Briefly, during anesthesia (2% isoflurane in pure O2) and with body temperature controlled by a heating pad (37°C), we removed the skin and connective tissue above the dorsal skull. To optically access auditory areas, we removed muscles above the respective skull location. After cleaning the skull, we applied a layer of UV-cure iBond followed by transparent dental cement (Tetric EvoFlow T1). Subsequently, we built a wall of dental cement “worms” (Charisma) the surrounding preparation and fixed a metal head post to the skull.

### Behavioral paradigms

Mice were trained in the auditory task only (n = 2), in the whisker-based tactile task only (n = 2), or in both tasks (n = 4). We designed both tasks as go/no-go discrimination tasks with delayed response. The delay period allowed a temporal separation of sensation period and reward-retrieval action as well as the study of short-term memory.

#### Tactile task

The behavioral setup and paradigm has been previously described (Chen et al., 2013; Gilad et al., 2018). After one second of baseline period, trials were initiated by a stimulus cue (2 beeps at 2 kHz, 100-ms duration with 50-ms interval) announcing the approaching texture (either a rough sandpaper of grit size P100; or a smooth sandpaper, P1200; pseudo-randomly presented with no more than 3 consecutive repetitions). The texture was presented to the right whisker pad, contralateral to the imaged hemisphere, and stayed in its final position for two seconds. Contacts between the whiskers and the texture typically occurred during this time window as well as up to about 1 second before texture stop (Gilad et al., 2018). Retraction of the texture triggered the start of the delay period (1 - 7 s). At the end of the delay period, a response cue (4 beeps at 4 kHz, 50-ms duration with 25-ms interval) signaled the start of the response window (2 seconds). A lick to the water spout in the response window was rewarded with a small drop of sweet water only in go-trials (‘Hit’). The spout was reachable at all times during trials. In no-go trials, incorrect licks were punished with white noise and a time out (~2 seconds; ‘false alarms’, FA). The absence of licks during the response window was neither rewarded nor punished in go (‘misses’) and no-go (‘correct-rejections’, CR) trials. Licks during the delay period (‘early licks’) were punished as in FA trials.

#### Auditory task

In order to compare cortical dynamics during sensation and short-term memory with another sensory modality, we designed an analogous task, in which mice had to discriminate two auditory tones (4 kHz, 8 kHz) in order to obtain reward. The loudspeaker was located on the right side of the head. In order to avoid shared sensory modality between cues and the relevant stimuli for discrimination, we exchanged the auditory ‘stimulus’ and ‘response’ cues of the tactile task with visual cues (single flash of 500-ms duration and 3 flashes of 150-ms duration at 100-ms interval, respectively). Trial structure and outcome remained untouched except of the timing between the stimulus cue and the sound onset, which randomly varied (2 ± 0.5 seconds).

#### Auditory task replay under anesthesia

In order to investigate the diverse A1 responses to auditory tones during the auditory task as well as to confirm the border between A1 and AD, we also replayed the auditory task with equal trial structure to expert mice during light anesthesia (1% isoflurane). In this case, only “go” trials were analyzed (**Figure S3**).

#### Training and performance

Three mice were conditioned to lick for the 4-kHz tone and 3 mice to lick for the 8-kHz tone. Of these 6 mice, 4 mice subsequently underwent additional training in the tactile discrimination task. From the mice conditioned to lick for the 4-kHz tone, one was conditioned to lick for the P100 texture, the other for the P1200 texture. The same was the case for the mice conditioned on the 8-kHz tone, so that all possible combinations were explored. Additionally, two more mice were trained on the tactile task, one with the P100 texture, the other with the P1200 texture serving as go stimulus. Performance was quantified as d-prime (d’ = *Z*(Hit/(Hit+Miss)) – *Z*(FA/(FA+CR)) (Chen et al., 2013), where *Z* denotes the inverse of the cumulative distribution function.

After recovery from surgery (5 to 7 days), mice were accustomed to the experimenter and head fixation. Water-scheduled mice were trained first to reliably lick to obtain a water reward. Next, they learned to report the go stimulus. At this stage, we gradually introduced the no-go stimulus. Once the mouse became an expert in discrimination (d>1.5), a short delay was introduced (hundreds of milliseconds). During this training phase, we successively prolonged the delay period based on the individual mouse’s performance (d’ and early lick rate). The complete training period typically lasted 3-10 weeks per task. Expert mice could reliably hold their decision until the start of the response window, while maintaining high performance (d’>1.5) and low percentage of early licks. If necessary, we granted additional training time so that mice would learn to not move (sit passively) for at least the first second of the delay period.

### Wide-field calcium imaging

In order to monitor simultaneously all areas in the dorsal cortex while an animal solved the task, we used the wide-field imaging approach. Excitation light emitted from a blue LED light (Thorlabs; M470L3) was filtered by the excitation filter (480/40 nm BrightLine HC), diffused, collimated, and directed to the left hemisphere of the mouse by a dichroic mirror (510 nm; AHF; Beamsplitter T510LPXRXT) filter cube (Thorlabs). The system comprises two objectives (Navitar; top objective, D-5095, 50 mm f0.95; bottom objective inverted, D-2595, 25 mm f0.95) with the dichroic mirror in between. Emission photons were collected through both objectives and the dichroic mirror, filtered (emission filter 514/30 nm, BrightLine HC), and recorded with a sensitive CMOS camera (Hamamatsu Orca Flash 4.0) mounted on top of the system. The field-of-view (~9 mm diameter) covered most of the dorsal cortex of the left hemisphere and part of the right hemisphere. Illumination power at the preparation was <0.1 mW/mm^2^. We record images of 512×512 pixels at 20 Hz frame rate. For these imaging conditions, we did not observe any photobleaching. At the beginning of each imaging day, we took a reference image of the skull and blood vessel pattern using a green fiber-coupled LED (Thorlabs).

#### Mapping and area selection

In order to align each individual brain to the Allen Mouse Common Coordinate Framework (Harris et al., 2019), we performed sensory mapping under light anesthesia (1% isoflurane, **Figure S1**). Contralateral to the imaging side, we presented five stimuli of different modalities: a loud speaker-coupled vibrating bar was used to stimulate whiskers, and forelimb and hindlimb paws (20 Hz for 2 s); white-noise, 4-kHz, and 8-kHz tones were applied for auditory stimulation (2-s duration); and a blue LED positioned in front of the eye provided a visual stimulation (100-ms duration; approximately at zero degree elevation and azimuth in the visual field). These set of stimuli yielded 5 functional spots -that together with anatomical landmarks (i.e. bregma, lamdba, and the midline) were used as anchoring points for registration of each individual brain to the atlas using a third-degree polynomial transformation. Using the atlas borders, we defined 25 areas of interest, with some minor manual modifications within these borders to fit the functional activity for each mouse (e.g., whiskers used by each mouse might differ). Other areas were defined by stereotaxic coordinates. Area definition and nomenclature: primary visual cortex (V1), Post-rhinal (POR), Posterior lateral (PL), Lateral intermediate (LI), Lateral medial (LM), Anterior lateral (AL), Rostrolateral (RL), Anterior (A), Anterior medial (AM), Posterior medial (PM), Retrosplenial dorsal (RD) and Retrosplenial angular (RA), Primary auditory (A1), Auditory dorsal (AD), auditory posterior and Temporal association area (Tea), Barrel cortex (BC; primary somatosensory whisker), Somatosensory nose (No), Somatosensory undetermined (UN), Somatosensory mouth (MO), Somatosensory forelimb (FL), Somatosensory hindlimb (HL), Somatosensory trunk (TR), Secondary somatosensory cortex (S2), M1, ALM (anterior lateral motor cortex; 2.5 anterior and 1.5 mm lateral from bregma (Li et al., 2015)) and secondary motor cortex (M2, 1.5 mm anterior and 0.5 mm lateral from bregma corresponding to Gilad et al., 2018). We defined here the areas at the posterior and lateral border of the visual cortex, mainly comprising P, POR, LM, and LI, as “posterolateral association” (PLA) areas.

We delineated the borders of AD-A1 and AD-S2 by a set of inclusion and exclusion criteria (**Figure S3D**). First, AD is the auditory area with highest activity and discrimination power during sensation for the auditory task (**Figure S2A**). Second, cortical activation during auditory mapping under anesthesia was mostly limited to A1 (**Figure S1**). Third, we observed the drastic difference in tone-evoked activation levels between AD and A1 during task performance but not during the replay of the auditory task under anesthesia or during the auditory cue of the tactile task (**Figure S2B, C)**. Four, during the sensation period of the tactile task, activity in S2 was higher than in AD (**Figure S2D**). After aligning each individual brain to the Allen Mouse Common Coordinate Framework, the area that fit with the criteria described above, localized best to the area denoted by the Allen Institute as AD.

### Body tracking

Simultaneous to wide-field imaging, we recorded movements of the body of the mouse during the task at 30 Hz (Body camera, The Imaging Source; DMK 22BUC03; 720×480 pixels). As imaging was performed in darkness, we illuminated the mouse using a 940-nm infrared LED.

#### Trial classification based on the body movements

We monitored major body movements of the mouse using a body camera. We focused on movements of the forelimb on the support pole accompanied by arching of the back. To extract a body movement vector, we combined the two regions of interest (ROIs), forelimbs and back. Specifically, movement was calculated as (1-corr(ft,ft+1)) where the correlation refers to the frame-to-frame correlation of these two ROIs. The movement vector was subsequently binarized by thresholding at 3 s.d. above baseline (defined as the 5^th^ percentile, ‘movement’ versus ‘quiet’). Irrespective of trial outcome (i.e. Hit, CR, etc.), individual trials were labeled as ‘active’ if the mouse moved at least 0.9 seconds during the sensation period (time window from −1.0 s to 2.0 s relative to sound onset/texture stop) or as ‘passive’ otherwise. Additionally, we calculate overall ‘activeness’ for each mouse as the percentage of active Hit trials. For delay period analyses, we only included active and passive trials, in which the mouse was quiet during the first second of the delay (starting 0.2 s after sound offset/texture retraction). In trials fitting these criteria, we truncated frames after the mouse movement onset. This restrictive analysis allowed us to largely exclude direct movement-related influences on cortical delay activity.

### Data analysis

We performed data analysis using Matlab software (Mathworks). 512×512 pixels images collected with the wide-field imaging setup were down-sampled to 256×256 pixels and pixels outside of the imaging area were discarded. We calculated ΔF/F by dividing fluorescence values for each trial and pixel by the average absolute fluorescence of several frames before the stimulus cue (baseline). To study neural dynamics related to sensation of the stimulus, we calculated baseline ΔF/F several frames before stimulus onset (sound onset or the earliest first touch of the whiskers on the texture reported by (Gilad et al., 2018) 1.1 s before texture stop, **Figure 2**). Next, trials were divided into the 5 categories of Hit, CR, FA, Miss, and Early licks. We focused our analyses on expert mice, thus only correct trials (Hit and CR) were considered.

#### Discrimination power between Hit and CR

To measure how well cortical areas (averaged ΔF/F over all the pixels included in a given area) or individual pixels could discriminate between Hit and CR trials, we calculated a receiver operating characteristics (ROC) curve and calculated its area under the curve (AUC).

#### Task and strategy indices

For these analyses, we focused on Hit trials from the four mice trained on both tasks and z-scored ΔF/F values in each trial. We defined indices, which describe how well areas can discriminate between behavioral strategies (strategy index *I*_S_) or between tasks (task index *I*_T_). For the strategy index, we performed for each pixel and time point an ROC curve for passive versus active trials and calculated *I*_S_ by adding the AUC values for auditory and tactile trials and adjusting to the index range of −1 (passive discrimination) to 1 (active discrimination; zero indicates no discrimination power):

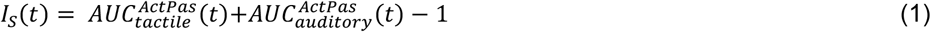

The task index was calculated in a similar way but taking into account that sensation periods were slightly different in the two tasks because the first-touch times varied in the tactile task (approximately −0.5 s before the texture reached its final position). Specifically, we aligned the peak time of the whisker-evoked average ΔF/F transient in BC to the peak time of the auditory-evoked ΔF/F signal in area A. Then we interpolated the segments of the ΔF/F transients in the time periods before and after the peak using the time points of cue and stimulus-end as fix points. While this alignment sharpened the task index values, we like to emphasize that the main results are also evident when using the non-aligned time courses. The task index was then calculated based on the ROC curves for tactile versus auditory trials, again adjusting *I*_T_ values to range from −1 (tactile) to 1 (auditory):

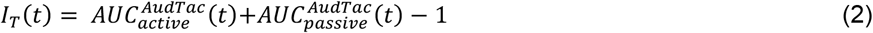

We obtained maps for the sensation and short-term memory periods for both indices by averaging index values for the respective time windows (0.1-0.2 s and 2.7-3.0 s). To plot index time courses, we calculated both indices for each brain area (using ΔF/F values averaged across all pixels within a given area) using equations 1 and 2. Then, we averaged the absolute values of each index across all cortical areas:

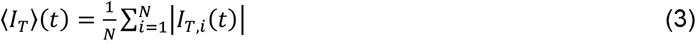

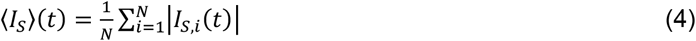

where *N* is the number of areas (here *N* = 26). In this case, index values ranged from 0 (no discrimination power) to 1 (maximum discrimination power for either task or strategy).

### Anatomy

#### Downloaded connectivity data from the Allen Institute

We downloaded and averaged all available experiments on transgenic lines for each relevant area from the Allen Institute (https://connectivity.brain-map.org/, Harris *et al*., 2019).

#### Retrograde labelling

In anaesthetized mice, we prepared small craniotomies over M2 and PLA areas. We injected in 3 mice AAV-retro-2-shortCAG-tdTomato-WPRE-SV40p(A) in M2 (1.65 mm anterior and 0.45 mm lateral from bregma) and AAV-retro-2-CAG-EGFP-WPRE-SV40p(A) in PLA areas (4.3 mm posterior and 3.45 mm lateral from bregma). In order to cover the entire cortical column, we injected each area with a minimum volume of 420 nl across cortical depth.

#### Hydrogel-based tissue clearing

The method used for hydrogel-based tissue clearing is described in detail elsewhere (Chung et al., 2013; Tomer et al., 2014; Yang et al., 2014; Ye et al., 2016). In short the brains were post-fixed for 48 hours in a Hydrogel solution (1% PFA, 4% Acrylamide, 0.05% Bis-Acrylamide) (Chung et al., 2013; Ye et al., 2016) before the hydrogel polymerization was induced at 37°C. Following the polymerization, the brains were immersed in 40mL of 8% SDS and kept shaking at room temperature until the tissue was cleared sufficiently (30 days). Finally, after 2-4 washes in PBS, the brains were put into a self-made refractive index matching solution (RIMS) (Yang et al., 2014) for the last clearing step. They were left to equilibrate in 5mL of RIMS for at least 4 days at RT before being imaged.

#### Cleared brain imaging

After clearing, brains were attached to a small weight and loaded into a 10 × 20 × 45 mm quartz cuvette (UQ-205, Portmann Instruments), then submerged in RIMS and imaged using a home-built mesoSPIM mesoscale single-plane illumination microscope (mesospim.org, Voigt et al. 2019). The sample cuvette was immersed in a 40 × 40 × 40 mm quartz cuvette (UQ-753, Portmann Instruments) filled with index-matching oil (19569, Code 50350, Cargille, nD=1.45) which allows sample XYZ & rotation movements without refocusing the detection path. The instrument consists of a dual-sided excitation path using a fiber-coupled multiline laser combiner (405, 488, 515, 561, 594, 647 nm, Omicron SOLE-6) and a detection path comprising an Olympus MVX-10 zoom macroscope with a 1× objective (Olympus MVPLAPO 1x), a filter wheel (Ludl 96A350), and a scientific CMOS (sCMOS) camera (Hamamatsu Orca Flash 4.0 V3). The excitation paths also contain galvo-scanners (Scanlab Dynaxis 3M 14-4) for light-sheet generation and reduction of streaking artifacts due to absorption of the light-sheet. In addition, the beam waist is scanned using electrically tunable lenses (ETL, Optotune EL-16-40-5D-TC-L) synchronized with the rolling shutter of the sCMOS camera. This axially scanned light-sheet mode (ASLM) leads to a uniform axial resolution across the field-of-view (FOV) of 4-10 μm (depending on zoom & wavelength). Image acquisition is done using custom software written in Python (https://github.com/mesoSPIM/mesoSPIM-control). We imaged with a field of view of 16.85 mm at 0.8× magnification (Pixel size: 8.23 μm) or 10.79 mm at 1.25× magnification (Pixel size: 5.27 μm). The laser/filter combinations were: EGFP: 488 nm excitation and a 520/35 bandpass filter (BrightLine HC, AHF); tdTomato: 561 nm excitation & 561 nm longpass (561LP Edge Basic, AHF); autofluorescence: 647 nm excitation & multiband emission filter (QuadLine Rejectionband ZET405/488/561/640, AHF).

#### Quantification of labelled neurons

Visualization and data analysis of 3D data were performed in Imaris 9.3 (Oxford instruments). BC, V1, RL and A were manually delineated using the expression pattern in the autofluorescence, green and red channels. Labelled neurons were automatically detected based on a combination of size (~35 μm) and intensity. Then, labelled neurons were manually confirmed and undetected ones manually marked.

#### Statistical analysis

In general, non-parametric two-tailed statistical tests were used, Wilcoxon signed rank test to compare the median between two populations. A one-way ANOVA was used when comparing all ROIs simultaneously or when comparing all time points of a time course.

## SUPPLEMENTARY INFORMATION

**Figure S1.**
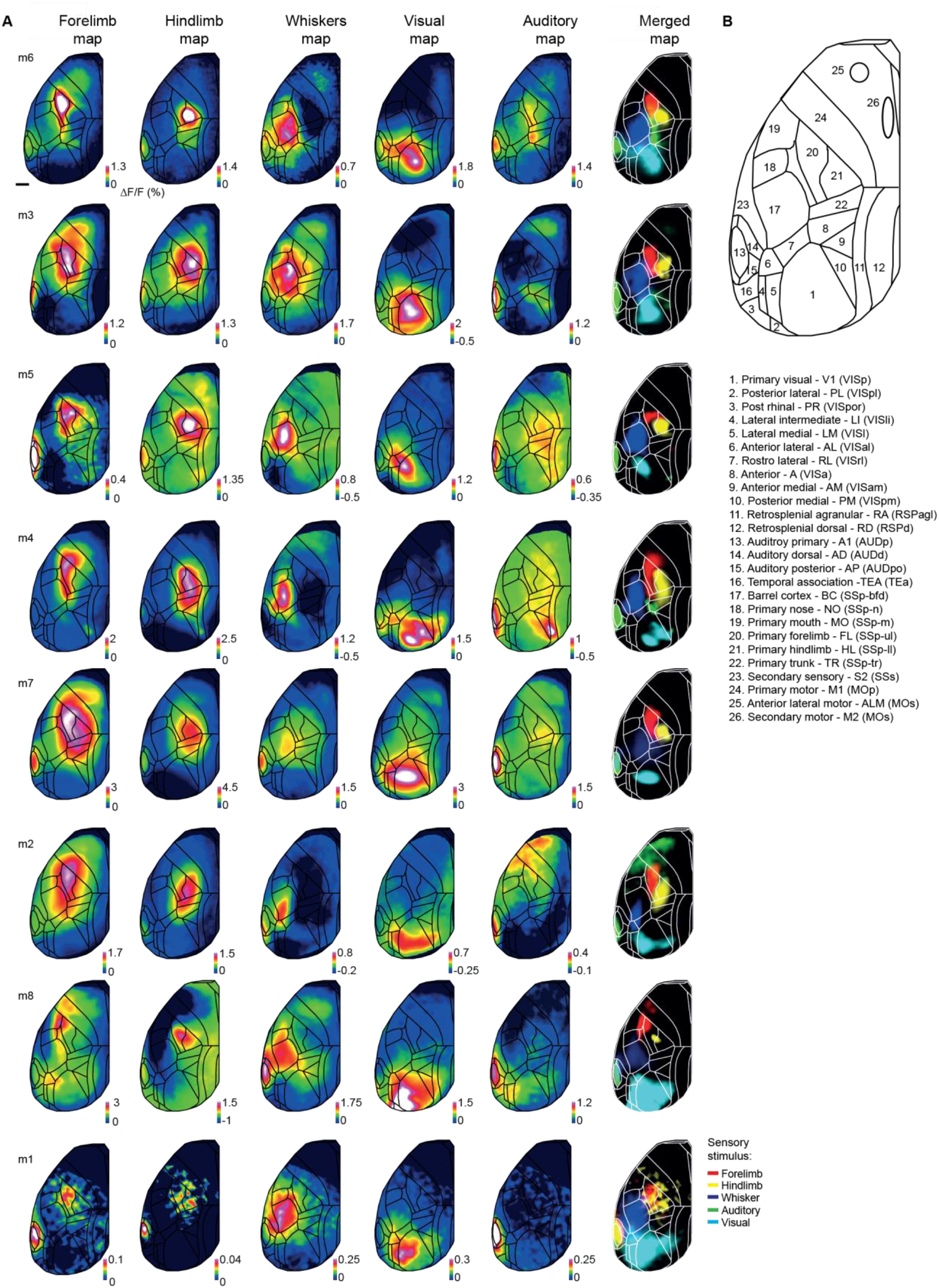
Related to Figure 2. Sensory mapping and alignment to Allen Brain Atlas coordinate framework for all mice. (A) Each mouse was presented with five stimuli of different modality: whisker, forelimb and hindlimb stimulation (somatosensory); white noise sound (auditory); and a blue LED flash (visual). Average stimulus-evoked ΔF/F maps were calculated and in the 6^th^ column all maps for an individual mouse are overlaid in different colors. These maps, together with bregma and lambda as anatomical landmarks, were used to align each brain to the Allen Mouse Common Coordinate Framework (area outline represents the top view of the atlas; all maps shown are already registered). Color denotes normalized fluorescence (ΔF/F). (B) Names and abbreviations of the 26 areas studied. For comparison, the abbreviations used by the Allen Institute are included in brackets.

**Figure S2.**
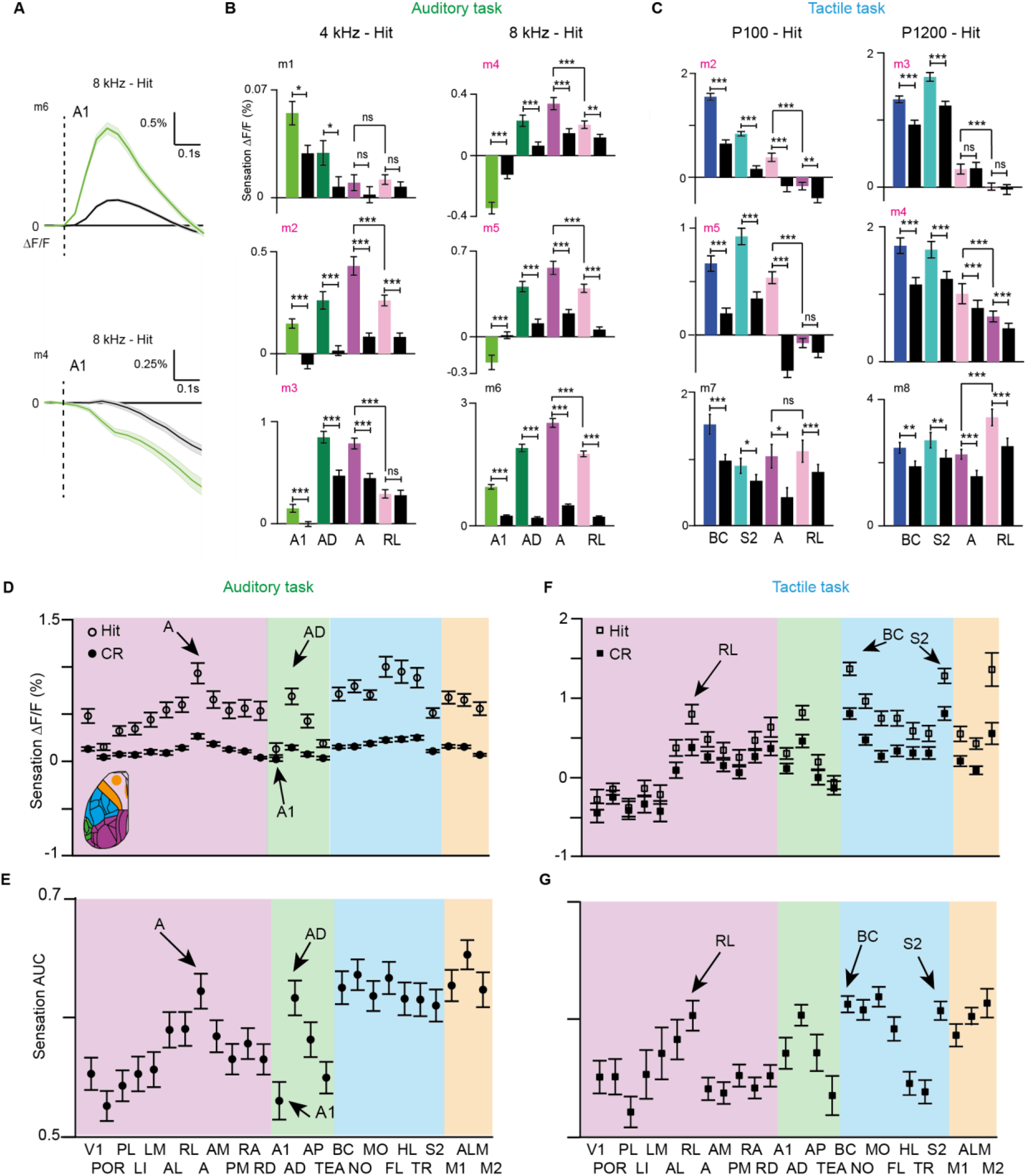
Related to Figure 2. Activity and decoding averages of individual mice and all areas during sensation. (**A**) Average A1 time course during sensation for two example mice (mouse 6 and 4). Note the different tone-evoked activity. (**B**) Average sensation activity for each mouse in Hit versus CR in A1, AD, A and RL in the auditory task; error bars are SEM across sessions. (**C**) Same as in (B) but for the tactile task. (**D**) Average sensation activity across mice in Hit versus CR in all dorsal cortical areas in the auditory task. Error bars are SEM across sessions (n=84) from 6 mice. (**E**) Average sensation area under the ROC curve (AUC) for Hit versus CR discrimination in all dorsal cortical areas in the auditory task. Error bars are SEM across sessions (n=84) from 6 mice. (**F**) Same as (D) but for the Tactile task. Error bars are SEM across sessions (n=78) from 6 mice. (**G**) Same as (E) but for the Tactile task. Error bars are SEM across sessions (n=88) from 6 mice. *p < 0.05, **p < 0.01, ***p < 0.001; n.s., not significant; Wilcoxon signed-rank test

**Figure S3.**
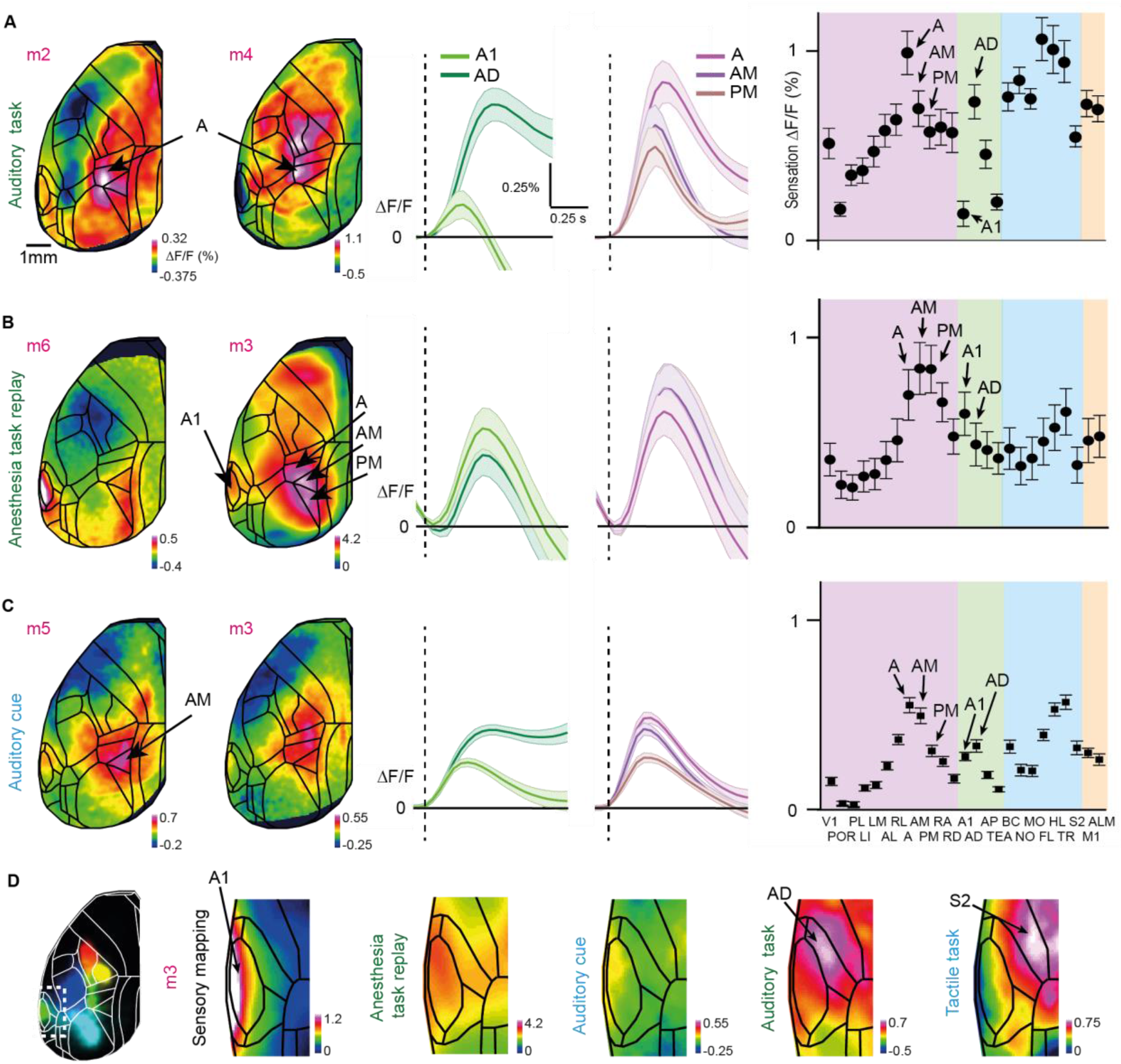
Related to Figure 2. A1-AD-S2 border definition and auditory-evoked activity in three different contexts: auditory discrimination task, task replay during anesthesia and texture-predicting auditory cue. (**A**) Tone-evoked activity during sensation in the auditory task. Session-averaged Hit sensation maps two example mice (left). Color scale bar indicates minimum and maximum percent ΔF/F. Average Hit sensation time course in A1 and AD; and A, AM and PM (middle). Average Hit sensation activity in in all dorsal cortical areas (right); error bars are SEM across sessions (n=84) from 6 mice. (**B**) Same as in (A) but for tone-evoked activity during anesthesia task replay. (**C**) Same as in (A) but for tone-evoked activity during auditory cue in the tactile task. (**D**) Definition of the A1-AD and AD-S2 borders in an example mice.

**Figure S4.**
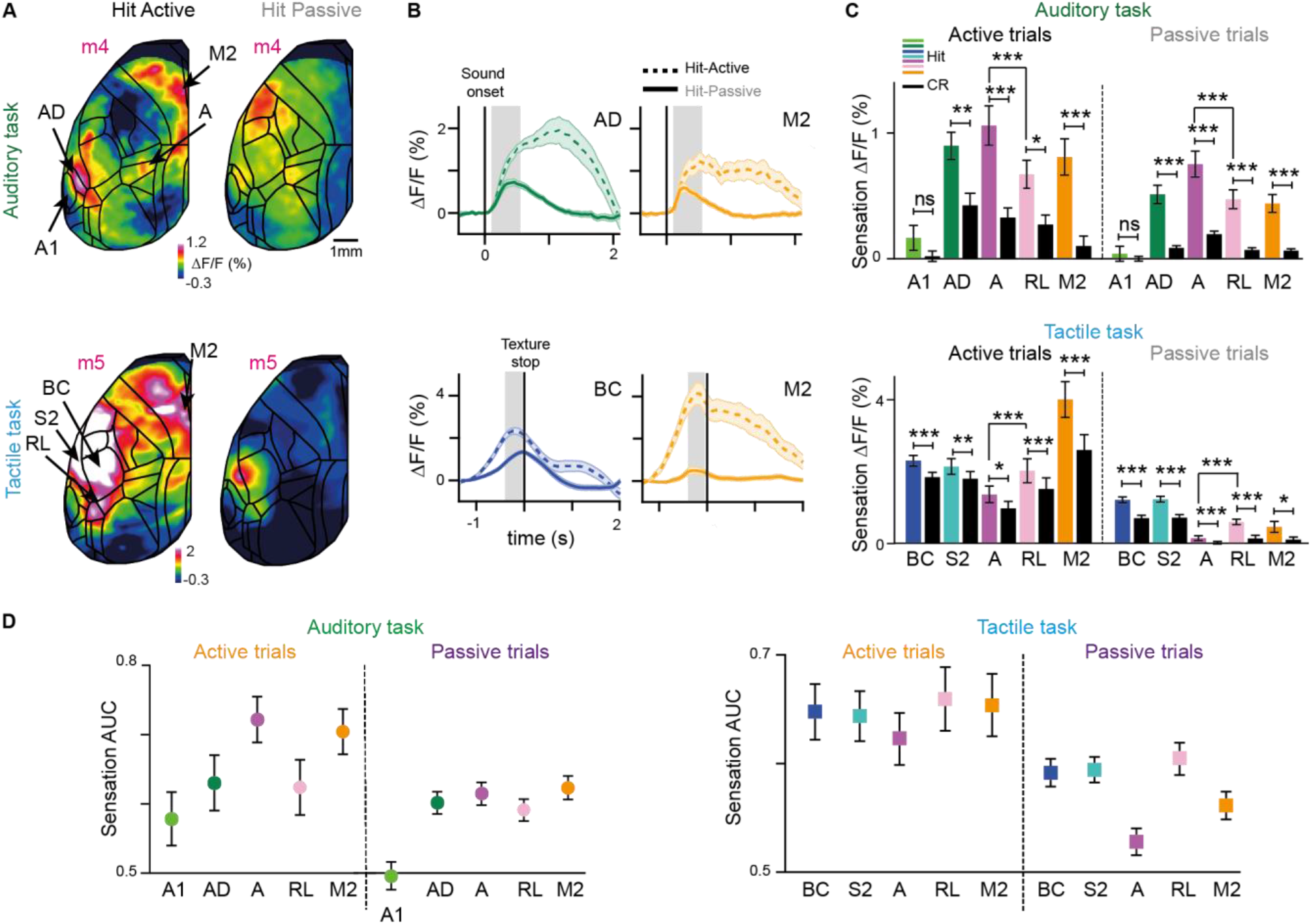
Related to Figure 2. Sensory-related cortical areas encode stimulus modality irrespective of strategy. (**A**) Top: session averaged sensation maps for an example mouse (m3) for Hit active (left) and Hit passive (right) from the auditory task. Color scale bar indicates ΔF/F percentage. Bottom: same as top, but for the tactile task. (**B**) Average time course Hit active (dashed line) versus Hit passive (solid line) in AD and M2 in the auditory (top); and in BC and M2 in the tactile (bottom) tasks. Error bars are SEM across sessions. (**C**) Top: average sensation activity in Hit active versus Hit passive in A1, AD, A and M2 in the auditory task. Bottom: same as top but for BC, S2, A and RL for the tactile task. Error bars are SEM across sessions (n=70, passive; n=28, active from 6 mice for the auditory task; and n=64, passive; n=30, active from 6 mice for the tactile task). (**D**) Hit/CR discrimination power, calculate as area under the ROC curve (AUC) in A1, AD, A, RL, and M2 in active and passive trials in the auditory task (left). Hit/CR discrimination power in BC, S2, A, RL, and M2 in the tactile task (right). Error bars are SEM across sessions (n=70 passive sessions from 5 mice and n=28 active sessions from 6 mice for the auditory task. For the tactile task, n=64 passive sessions from 4 mice; n=30 active sessions from 5 mice, for the tactile task). *p < 0.05, **p < 0.01, ***p < 0.001; n.s., not significant; Wilcoxon signed-rank test.

**Figure S5.**
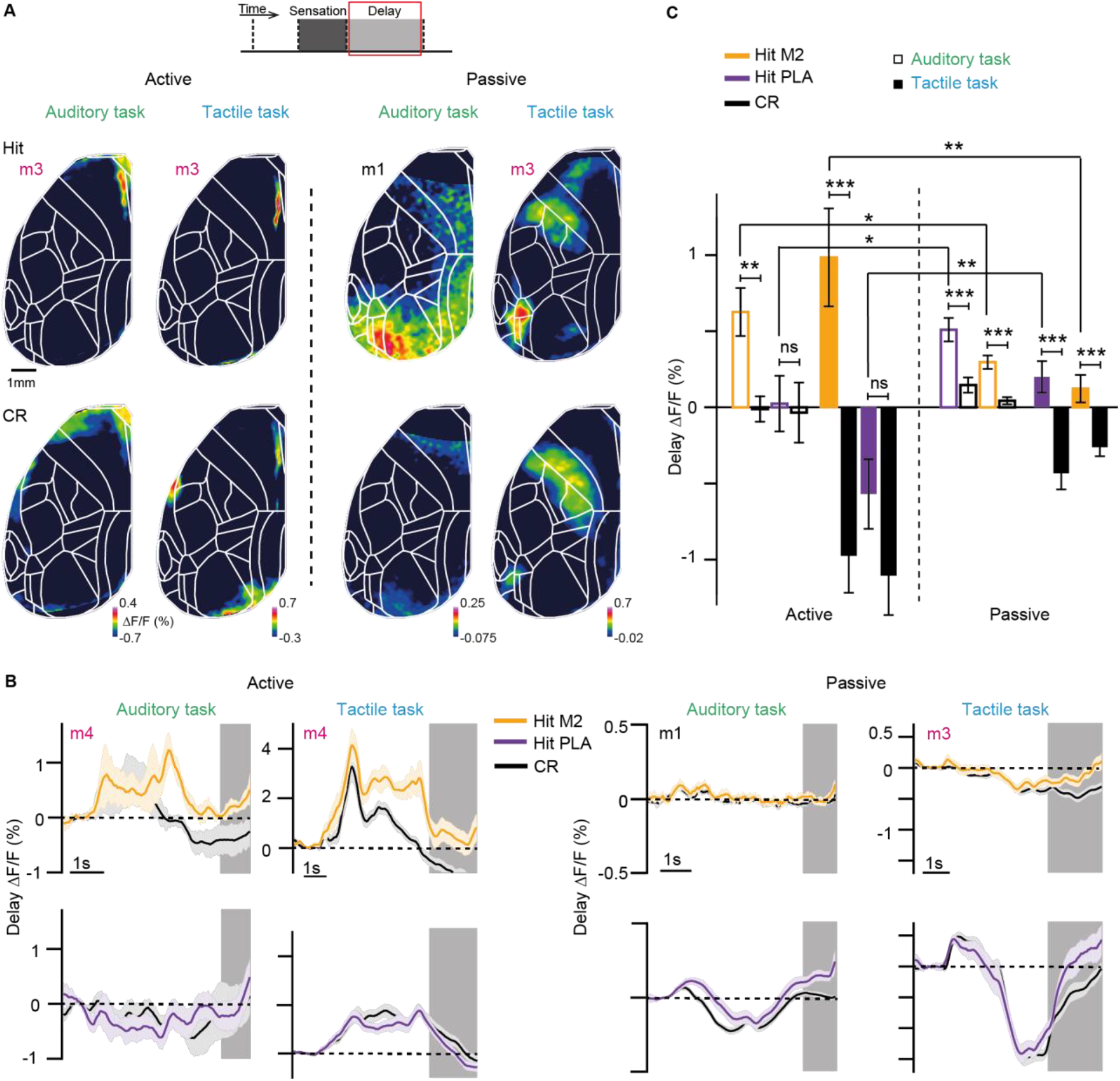
Related to Figure 4. Delay activity maps, time courses and average delay activity in both strategies and tasks. (**A**) Session averaged delay maps for active (left) and passive (right) Hits and CRs trials for two example mice in both tasks. Mouse 3 was trained in both tasks; mouse 1 only in the auditory. Color scale bar indicates ΔF/F percentage. (**B**) Average time course Hit versus CR for both tasks and strategies in M2 and P. Error bars are SEM across sessions. Grey indicates delay period. (**C**) Average delay activity in Hit versus CR in M2 and P for both strategies and tasks. Error bars are SEM across sessions. Auditory task: passive sessions n=70 from 6 mice; active sessions n= 28 from 4 mice. Tactile task: passive sessions n=64 from 4 mice; active sessions n= 30 from 5 mice. *p < 0.05, **p < 0.01, ***p < 0.001; n.s., not significant; Wilcoxon signed-rank test

**Figure S6.**
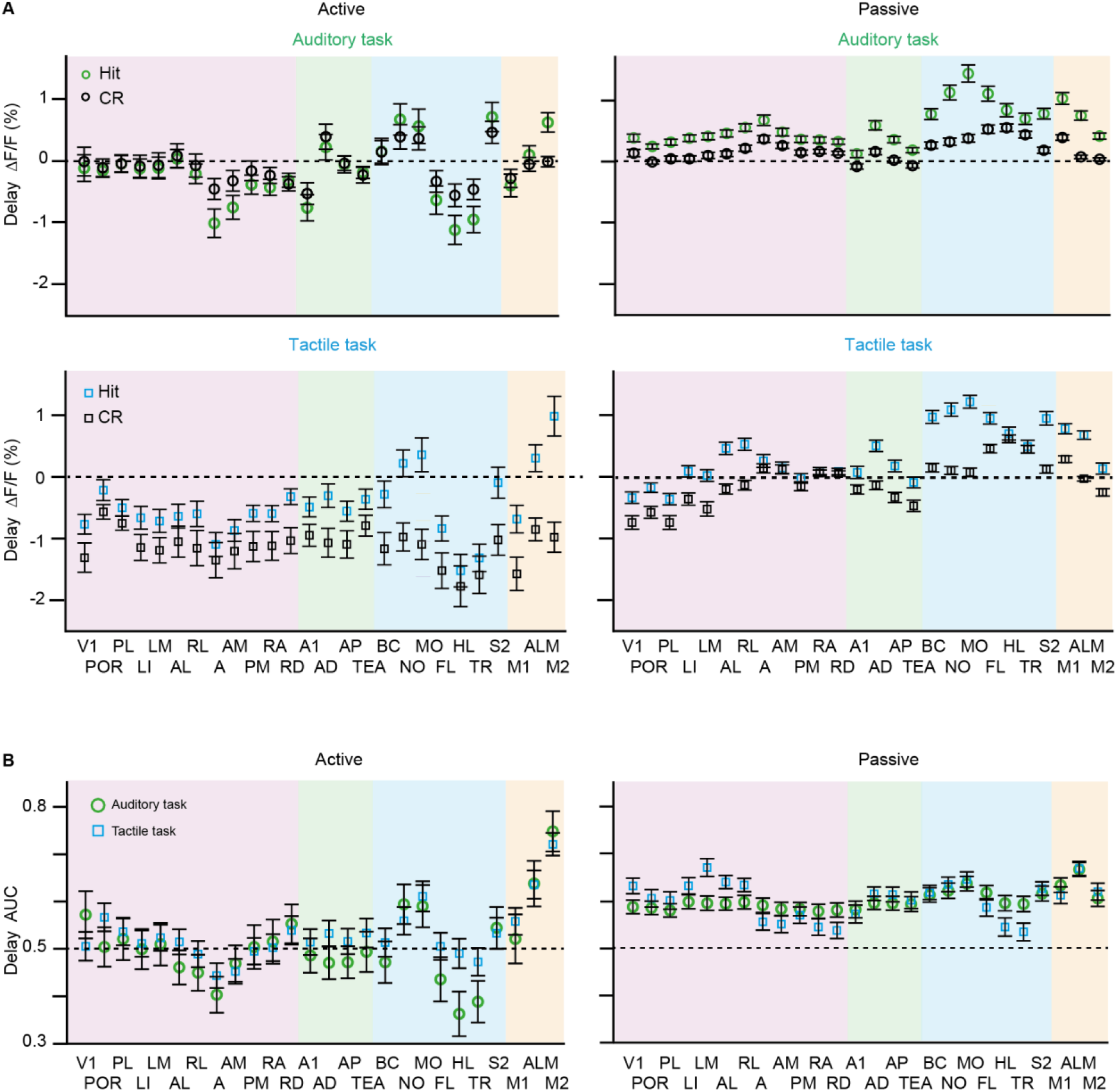
Related to Figure 4. Average delay activity in Hit versus CR and delay AUC for Hit versus CR discrimination in all areas of the dorsal neocortex. (**A**) Average delay activity in Hit versus CR in all areas of the dorsal neocortex for both strategies and tasks. Error bars are SEM across sessions. Auditory task: passive sessions n=70 from 6 mice; active sessions n= 28 from 4 mice. Tactile task: passive sessions n=64 from 4 mice; active sessions n= 30 from 5 mice. (**B**) Same as (A) but for AUC for Hit versus CR discrimination.

**Figure S7.**
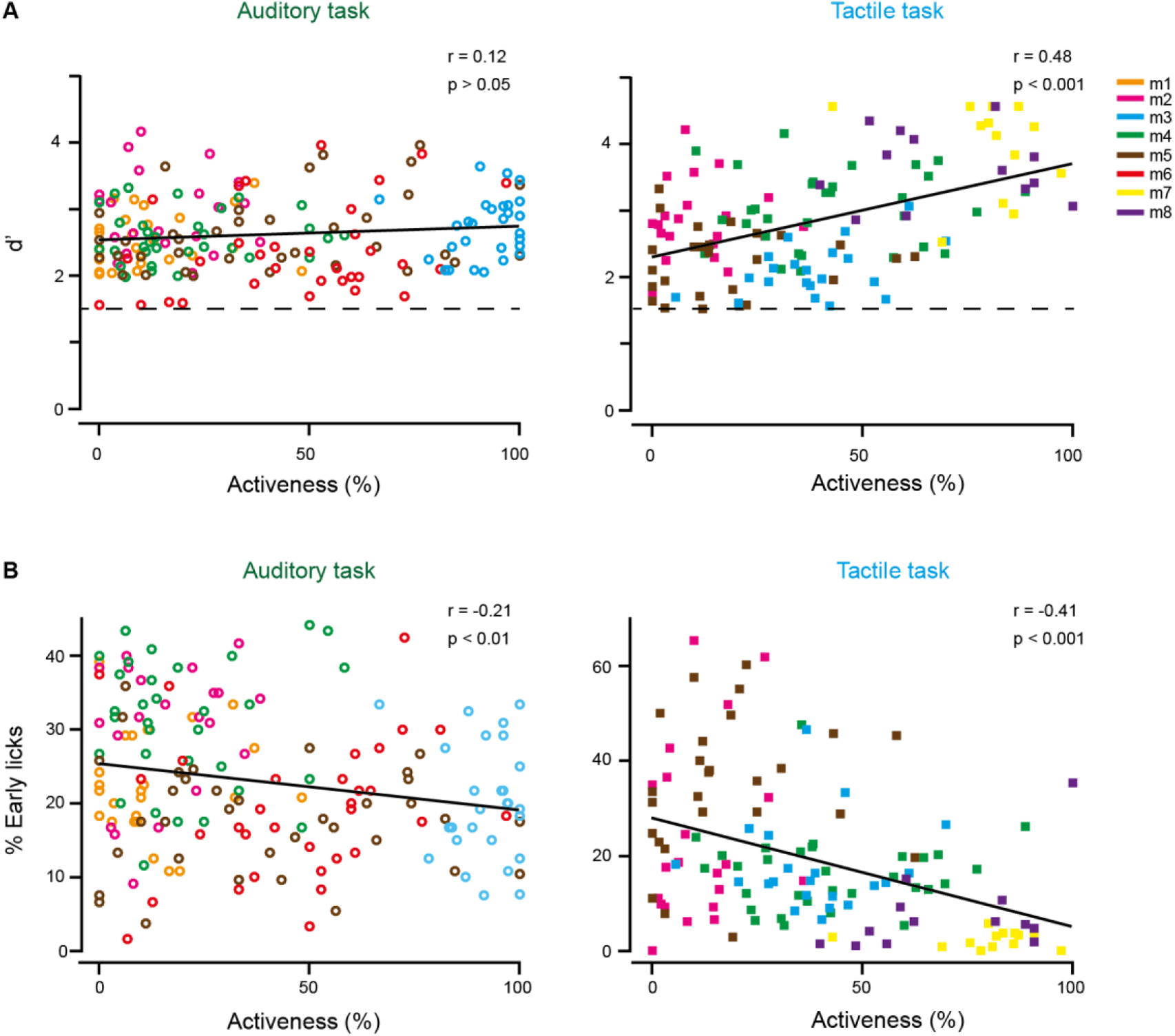
Correlations between performance and activeness. (**A**) Correlation between performance (measured as d’) and activeness for the auditory task (left) and the tactile task (right). Dashed line indicates expert threshold at d’ = 1.5. (**B**) Correlation between the percentage of early licks and activeness for the auditory task (left) and the tactile task (right). Data points are individual sessions from the different mice.

## REFERENCES

Allen, W.E., Kauvar, I. V., Chen, M.Z., Richman, E.B., Yang, S.J., Chan, K., Gradinaru, V., Deverman, B.E., Luo, L., and Deisseroth, K. (2017). Global representations of goal-directed behavior in distinct cell types of mouse neocortex. Neuron 94, 891–907.e6.

Amedi, A., Von Kriegstein, K., Van Atteveldt, N.M., Beauchamp, M.S., and Naumer, M.J. (2005). Functional imaging of human crossmodal identification and object recognition. In Experimental Brain Research 166, 559–571.

Atiani, S., David, S. V., Elgueda, D., Locastro, M., Radtke-Schuller, S., Shamma, S.A., and Fritz, J.B. (2014). Emergent selectivity for task-relevant stimuli in higher-order auditory cortex. Neuron 82, 486–499.

Barthas, F., and Kwan, A.C. (2017). Secondary motor cortex: Where ‘sensory’ meets ‘motor’ in the rodent frontal cortex. Trends Neurosci. 40, 181–193.

Beauchamp, M.S. (2005). See me, hear me, touch me: Multisensory integration in lateral occipital-temporal cortex. Curr. Opin. Neurobiol. 15, 145–153.

Chen, J.L., Carta, S., Soldado-Magraner, J., Schneider, B.L., and Helmchen, F. (2013). Behaviour-dependent recruitment of long-range projection neurons in somatosensory cortex. Nature 499, 336–340.

Chen, T.W., Li, N., Daie, K., and Svoboda, K. (2017). A map of anticipatory activity in mouse motor cortex. Neuron 94, 866–879.e4.

La Chioma, A., Bonhoeffer, T., and Hübener, M. (2019). Area-specific mapping of binocular disparity across mouse visual cortex. Curr. Biol. 29, 2954–2960.e5.

Chung, K., Wallace, J., Kim, S.Y., Kalyanasundaram, S., Andalman, A.S., Davidson, T.J., Mirzabekov, J.J., Zalocusky, K.A., Mattis, J., Denisin, A.K., et al. (2013). Structural and molecular interrogation of intact biological systems. Nature 497, 332–337.

Clancy, K.B., Orsolic, I., and Mrsic-Flogel, T.D. (2019). Locomotion-dependent remapping of distributed cortical networks. Nat. Neurosci. 22, 778–786.

Colombo, M., D’Amato, M.R., Rodman, H.R., and Gross, C.G. (1990). Auditory association cortex lesions impair auditory short-term memory in monkeys. Science 247, 336–338.

Dhawale, A.K., Miyamoto, Y.R., Smith, M.A., and Ölveczky, B.P. (2019). Adaptive regulation of motor variability. Curr. Biol. 29, 3551–3562.e7.

Dong, C., Qin, L., Zhao, Z., Zhong, R., and Sato, Y. (2013). Behavioral modulation of neural encoding of click-trains in the primary and nonprimary auditory cortex of cats. J. Neurosci. 33, 13126–13137.

Elgueda, D., Duque, D., Radtke-Schuller, S., Yin, P., David, S. V., Shamma, S.A., and Fritz, J.B. (2019). State-dependent encoding of sound and behavioral meaning in a tertiary region of the ferret auditory cortex. Nat. Neurosci. 22, 447–459.

Esmaeili, V., and Diamond, M.E. (2019). Neuronal correlates of tactile working memory in prefrontal and vibrissal somatosensory cortex. Cell Rep. 27, 3167–3181.e5.

Francis, N.A., Winkowski, D.E., Sheikhattar, A., Armengol, K., Babadi, B., and Kanold, P.O. (2018). Small networks encode decision-making in primary auditory cortex. Neuron 97, 885–897.e6.

Gilad, A., Gallero-Salas, Y., Groos, D., and Helmchen, F. (2018). Behavioral strategy determines frontal or posterior location of short-term memory in neocortex. Neuron 99, 814–828.e7.

Gimenez, T.L., Lorenc, M., and Jaramillo, S. (2015). Adaptive categorization of sound frequency does not require the auditory cortex in rats. J. Neurophysiol. 114, 1137–1145.

Glickfeld, L.L., and Olsen, S.R. (2017). Higher-order areas of the mouse visual cortex. Annu. Rev. Vis. Sci. 3, 251–273.

Goard, M.J., Pho, G.N., Woodson, J., and Sur, M. (2016). Distinct roles of visual, parietal, and frontal motor cortices in memory-guided sensorimotor decisions. Elife 5, 1–30.

Guo, Z. V., Li, N., Huber, D., Ophir, E., Gutnisky, D., Ting, J.T., Feng, G., and Svoboda, K. (2014). Flow of cortical activity underlying a tactile decision in mice. Neuron 81, 179–194.

Harris, J.A., Mihalas, S., Hirokawa, K.E., Whitesell, J.D., Choi, H., Bernard, A., Bohn, P., Caldejon, S., Casal, L., Cho, A., et al. (2019). Hierarchical organization of cortical and thalamic connectivity. Nature 575, 195–202.

Harvey, C.D., Coen, P., and Tank, D.W. (2012). Choice-specific sequences in parietal cortex during a virtual-navigation decision task. Nature 484, 62–68.

Hovde, K., Gianatti, M., Witter, M.P., and Whitlock, J.R. (2019). Architecture and organization of mouse posterior parietal cortex relative to extrastriate areas. Eur. J. Neurosci. 49, 1313–1329.

Inagaki, H.K., Inagaki, M., Romani, S., and Svoboda, K. (2018). Low-dimensional and monotonic preparatory activity in mouse anterior lateral motor cortex. J. Neurosci. 38, 4163–4185.

Jaramillo, S., and Zador, A.M. (2011). The auditory cortex mediates the perceptual effects of acoustic temporal expectation. Nat. Neurosci. 14, 246–253.

Kamigaki, T., and Dan, Y. (2017). Delay activity of specific prefrontal interneuron subtypes modulates memory-guided behavior. Nat. Neurosci. 20, 854–863.

Kato, H.K., Gillet, S.N., and Isaacson, J.S. (2015). Flexible sensory representations in auditory cortex driven by behavioral relevance. Neuron 88, 1027–1039.

Kuchibhotla, K. V., Gill, J. V., Lindsay, G.W., Papadoyannis, E.S., Field, R.E., Sten, T.A.H., Miller, K.D., and Froemke, R.C. (2017). Parallel processing by cortical inhibition enables context-dependent behavior. Nat. Neurosci. 20, 62–71.

Kuroki, S., Yoshida, T., Tsutsui, H., Iwama, M., Ando, R., Michikawa, T., Miyawaki, A., Ohshima, T., and Itohara, S. (2018). Excitatory neuronal hubs configure multisensory integration of slow waves in association cortex. Cell Rep. 22, 2873–2885.

Lee, S.H., Kravitz, D.J., and Baker, C.I. (2013). Goal-dependent dissociation of visual and prefrontal cortices during working memory. Nat. Neurosci. 16, 997–999.

Lee, T., Alloway, K.D., and Kim, U. (2011). Interconnected cortical networks between primary somatosensory cortex septal columns and posterior parietal cortex in rat. J. Comp. Neurol. 519, 405–419.

Li, N., Chen, T.W., Guo, Z. V., Gerfen, C.R., and Svoboda, K. (2015). A motor cortex circuit for motor planning and movement. Nature 519, 51–56.

Lippert, M.T., Takagaki, K., Kayser, C., and Ohl, F.W. (2013). Asymmetric multisensory interactions of visual and somatosensory responses in a region of the rat parietal cortex. PLoS One 8 (5), e63631.

Liu, D., Gu, X., Zhu, J., Zhang, X., Han, Z., Yan, W., Cheng, Q., Hao, J., Fan, H., Hou, R., et al. (2014). Medial prefrontal activity during delay period contributes to learning of a working memory task. Science. 346, 458–463.

Lucan, J.N., Foxe, J.J., Gomez-Ramirez, M., Sathian, K., and Molholm, S. (2010). Tactile shape discrimination recruits human lateral occipital complex during early perceptual processing. Hum. Brain Mapp. 31, 1813–1821.

Lyamzin, D., and Benucci, A. (2019). The mouse posterior parietal cortex: Anatomy and functions. Neurosci. Res. 140, 14–22.

Madisen, L., Garner, A.R., Shimaoka, D., Chuong, A.S., Klapoetke, N.C., Li, L., van der Bourg, A., Niino, Y., Egolf, L., Monetti, C., et al. (2015). Transgenic mice for intersectional targeting of neural sensors and effectors with high specificity and performance. Neuron 85, 942–958.

Makino, H., Ren, C., Liu, H., Kim, A.N., Kondapaneni, N., Liu, X., Kuzum, D., and Komiyama, T. (2017). Transformation of cortex-wide emergent properties during motor learning. Neuron 94, 880–890.e8.

Le Merre, P., Esmaeili, V., Charrière, E., Galan, K., Salin, P.A., Petersen, C.C.H., and Crochet, S. (2018). Reward-based learning drives rapid sensory signals in medial prefrontal cortex and dorsal hippocampus necessary for goal-directed behavior. Neuron 97, 83–91.e5.

Mohajerani, M.H., Chan, A.W., Mohsenvand, M., Ledue, J., Liu, R., McVea, D.A., Boyd, J.D., Wang, Y.T., Reimers, M., and Murphy, T.H. (2013). Spontaneous cortical activity alternates between motifs defined by regional axonal projections. Nat. Neurosci. 16, 1426–1435.

Mohan, H., Gallero-Salas, Y., Carta, S., Sacramento, J., Laurenczy, B., Sumanovski, L.T., De Kock, C.P.J., Helmchen, F., and Sachidhanandam, S. (2018a). Sensory representation of an auditory cued tactile stimulus in the posterior parietal cortex of the mouse. Sci. Rep. 8, 1–13.

Mohan, H., de Haan, R., Mansvelder, H.D., and de Kock, C.P.J. (2018b). The posterior parietal cortex as integrative hub for whisker sensorimotor information. Neuroscience 368, 240–245.

Mohan, H., de Haan, R., Broersen, R., Pieneman, A.W., Helmchen, F., Staiger, J.F., Mansvelder, H.D., and de Kock, C.P.J. (2019). Functional architecture and encoding of tactile sensorimotor behavior in rat posterior parietal cortex. J. Neurosci. 39, 7332–7343.

Morcos, A.S., and Harvey, C.D. (2016). History-dependent variability in population dynamics during evidence accumulation in cortex. Nat. Neurosci. 19, 1672–1681.

Murakami, M., Vicente, M.I., Costa, G.M., and Mainen, Z.F. (2014). Neural antecedents of self-initiated actions in secondary motor cortex. Nat. Neurosci. 17, 1574–1582.

Musall, S., Kaufman, M.T., Juavinett, A.L., Gluf, S., and Churchland, A.K. (2019). Single-trial neural dynamics are dominated by richly varied movements. Nat. Neurosci. 22, 1677–1686.

Nikbakht, N., Tafreshiha, A., Zoccolan, D., and Diamond, M.E. (2018). Supralinear and supramodal integration of visual and tactile signals in rats: Psychophysics and neuronal mechanisms. Neuron 97, 626–639.e8.

Niwa, M., Johnson, J.S., O’Connor, K.N., and Sutter, M.L. (2013). Differences between primary auditory cortex and auditory belt related to encoding and choice for AM sounds. J. Neurosci. 33, 8378–8395.

Odoemene, O., Pisupati, S., Nguyen, H., and Churchland, A.K. (2018). Visual evidence accumulation guides decision-making in unrestrained mice. J. Neurosci. 38, 10143–10155.

Oh, S.W., Harris, J.A., Ng, L., Winslow, B., Cain, N., Mihalas, S., Wang, Q., Lau, C., Kuan, L., Henry, A.M., et al. (2014). A mesoscale connectome of the mouse brain. Nature 508, 207–214.

Ohl, F.W., Wetzel, W., Wagner, T., Rech, A., and Scheich, H. (1999). Bilateral ablation of auditory cortex in Mongolian gerbil affects discrimination of frequency modulated tones but not of pure tones. Learn. Mem. 6, 347–362.

Olcese, U., Iurilli, G., and Medini, P. (2013). Cellular and synaptic architecture of multisensory integration in the mouse neocortex. Neuron 79, 579–593.

Otazu, G.H., Tai, L.H., Yang, Y., and Zador, A.M. (2009). Engaging in an auditory task suppresses responses in auditory cortex. Nat. Neurosci. 12, 646–654.

Pho, G.N., Goard, M.J., Woodson, J., Crawford, B., and Sur, M. (2018). Task-dependent representations of stimulus and choice in mouse parietal cortex. Nat. Commun. 9, 2596.

Pinto, L., Rajan, K., DePasquale, B., Thiberge, S.Y., Tank, D.W., and Brody, C.D. (2019). Task-dependent changes in the large-scale dynamics and necessity of cortical regions. Neuron 104, 810–824.e9.

Quak, M., London, R.E., and Talsma, D. (2015). A multisensory perspective of working memory. Front. Hum. Neurosci. 9, 1–11.

Ramesh, R.N., Burgess, C.R., Sugden, A.U., Gyetvan, M., and Andermann, M.L. (2018). Intermingled ensembles in visual association cortex encode stimulus identity or predicted outcome. Neuron 100, 900–915.e9.

Sakurai, Y. (1994). Involvement of auditory cortical and hippocampal neurons in auditory working memory and reference memory in the rat. J. Neurosci. 14, 2606–2623.

Salkoff, D.B., Zagha, E., McCarthy, E., and McCormick, D.A. (2019). Movement and performance explain widespread cortical activity in a visual detection task. Cereb. Cortex, bhz206.

Schroeder, C.E., Wilson, D.A., Radman, T., Scharfman, H., and Lakatos, P. (2010). Dynamics of active sensing and perceptual selection. Curr. Opin. Neurobiol. 20, 172–176.

Shuler, M.G., and Bear, M.F. (2006). Reward timing in the primary visual cortex. Science 311, 1606–1609.

Siegel, M., Buschman, T.J., and Miller, E.K. (2015). Cortical information flow during flexible sensorimotor decisions. Science. 348, 1352–1355.

Silasi, G., Xiao, D., Vanni, M.P., Chen, A.C.N., and Murphy, T.H. (2016). Intact skull chronic windows for mesoscopic wide-field imaging in awake mice. J. Neurosci. Methods 267, 141–149.

Sreenivasan, K.K., and D’Esposito, M. (2019). The what, where and how of delay activity. Nat. Rev. Neurosci. 20, 466–481.

Stringer, C., Pachitariu, M., Steinmetz, N., Reddy, C.B., Carandini, M., and Harris, K.D. (2019). Spontaneous behaviors drive multidimensional, brainwide activity. Science 364, 255.

Sul, J.H., Jo, S., Lee, D., and Jung, M.W. (2011). Role of rodent secondary motor cortex in value-based action selection. Nat. Neurosci. 14, 1202–1210.

Talwar, S.K., Musial, P.G., and Gerstein, G.L. (2001). Role of mammalian auditory cortex in the perception of elementary sound properties. J. Neurophysiol. 85, 2350–2358.

Tomer, R., Ye, L., Hsueh, B., and Deisseroth, K. (2014). Advanced CLARITY for rapid and high-resolution imaging of intact tissues. Nat. Protoc. 9, 1682–1697.

Tsunada, J., Liu, A.S.K., Gold, J.I., and Cohen, Y.E. (2015). Causal contribution of primate auditory cortex to auditory perceptual decision-making. Nat. Neurosci. 19, 135–142.

Vanni, M.P., and Murphy, T.H. (2014). Mesoscale transcranial spontaneous activity mapping in GCaMP3 transgenic mice reveals extensive reciprocal connections between areas of somatomotor cortex. J. Neurosci. 34, 15931–15946.

Voigt, F.F., Kirschenbaum, D., Platonova, E., Pagès, S., Campbell, R.A.A., Kastli, R., Schaettin, M., Egolf, L., van der Bourg, A., Bethge, P., et al. (2019). The mesoSPIM initiative: open-source light-sheet microscopes for imaging cleared tissue. Nat. Methods. 16, 1105–1108

Wang, Q., Sporns, O., and Burkhalter, A. (2012). Network analysis of corticocortical connections reveals ventral and dorsal processing streams in mouse visual cortex. J. Neurosci. 32, 4386–4399.

Wekselblatt, J.B., Flister, E.D., Piscopo, D.M., and Niell, C.M. (2016). Large-scale imaging of cortical dynamics during sensory perception and behavior. J. Neurophysiol. 115, 2852–2866.

Wilber, A.A., Clark, B.J., Clark, B.J., Mesina, L., Vos, J.M., and McNaughton, B.L. (2015). Cortical connectivity maps reveal anatomically distinct areas in the parietal cortex of the rat. Front. Neural Circuits 8, 1–15.

Xin, Y., Zhong, L., Zhang, Y., Zhou, T., Pan, J., and Xu, N. long (2019). Sensory-to-category transformation via dynamic reorganization of ensemble structures in mouse auditory cortex. Neuron 103, 909–921.e6.

Yang, B., Treweek, J.B., Kulkarni, R.P., Deverman, B.E., Chen, C.K., Lubeck, E., Shah, S., Cai, L., and Gradinaru, V. (2014). Single-cell phenotyping within transparent intact tissue through whole-body clearing. Cell 158, 945–958.

Ye, L., Allen, W.E., Thompson, K.R., Tian, Q., Hsueh, B., Ramakrishnan, C., Wang, A.C., Jennings, J.H., Adhikari, A., Halpern, C.H., et al. (2016). Wiring and molecular features of prefrontal ensembles representing distinct experiences. Cell 165, 1776–1788.

Zhong, L., Zhang, Y., Duan, C.A., Deng, J., Pan, J., and Xu, N. long (2019). Causal contributions of parietal cortex to perceptual decision-making during stimulus categorization. Nat. Neurosci. 22, 963–973.

Zingg, B., Hintiryan, H., Gou, L., Song, M.Y., Bay, M., Bienkowski, M.S., Foster, N.N., Yamashita, S., Bowman, I., Toga, A.W., et al. (2014). Neural networks of the mouse neocortex. Cell 156, 1096–1111.

